# Loss of neuronal lysosomal acid lipase drives amyloid pathology in Alzheimer’s disease

**DOI:** 10.1101/2024.06.09.596693

**Authors:** Alexandra M Barnett, Lamar Dawkins, Jian Zou, Elizabeth McNair, Viktoriya D Nikolova, Sheryl S Moy, Greg T Sutherland, Julia Stevens, Meagan Colie, Kemi Katemboh, Hope Kellner, Corina Damian, Sagan DeCastro, Ryan P Vetreno, Leon G Coleman

## Abstract

Underlying drivers of late-onset Alzheimer’s disease (LOAD) pathology remain unknown. However, multiple biologically diverse risk factors share a common pathological progression. To identify convergent molecular abnormalities that drive LOAD pathogenesis we compared two common midlife risk factors for LOAD, heavy alcohol use and obesity. This revealed that disrupted lipophagy is an underlying cause of LOAD pathogenesis. Both exposures reduced lysosomal flux, with a loss of neuronal lysosomal acid lipase (LAL). This resulted in neuronal lysosomal lipid (NLL) accumulation, which opposed Aβ localization to lysosomes. Neuronal LAL loss both preceded (with aging) and promoted (targeted knockdown) Aβ pathology and cognitive deficits in AD mice. The addition of recombinant LAL *ex vivo* and neuronal LAL overexpression *in vivo* prevented amyloid increases and improved cognition. In WT mice, neuronal LAL declined with aging and correlated negatively with entorhinal Aβ. In healthy human brain, LAL also declined with age, suggesting this contributes to the age-related vulnerability for AD. In human LOAD LAL was further reduced, correlated negatively with Aβ_1-42_, and occurred with polymerase pausing at the LAL gene. Together, this finds that the loss of neuronal LAL promotes NLL accumulation to impede degradation of Aβ in neuronal lysosomes to drive AD amyloid pathology.

**Summary:** Cellular and molecular drivers of late-onset Alzheimer’s disease (LOAD) are unknown, though several risk factors account for the majority of disease incidence^1–5^. Though diverse in their biological natures, each of these risk exposures converge on a shared pathological progression with the accumulation of amyloid early in the disease. Human genetic and transcriptomic studies suggest a role for altered lipid metabolism^6–9^, though the mechanism has been unknown. Here, using two common midlife risk exposures for LOAD, we found that dysfunctional lipophagy caused by the loss of lysosomal acid lipase (LAL) promotes early LOAD pathogenesis. Both midlife obesity and heavy alcohol reduced neuronal LAL, causing an increase in neuronal lysosomal lipid, and a subsequent accumulation of Aβ in the extra-lysosomal cytosol. This loss of LAL preceded and promoted Aβ pathology and cognitive deficits in AD mice. The addition of recombinant LAL *ex vivo* and neuronal LAL overexpression *in vivo* prevented increases in amyloid and improved cognition. In human brain, LAL declined with age in healthy subjects, similar to rodents, showing robust losses in LOAD subjects with polymerase pausing. Together, this implicates neuronal LAL loss in LOAD pathogenesis and presents LAL as a promising diagnostic, preventative, and/or therapeutic target for AD.

## Main Text

Alzheimer’s disease (AD) is the most common form of dementia with over 90% of cases being sporadic/late-onset (LOAD). Though the etiology of LOAD is often unknown, risk factors such as heavy smoking or alcohol use, diabetes, hypertension, and obesity account for majority of risk^1–5^. Though these exposures have diverse biological consequences, they result in the same pathological progression – accumulation of intraneuronal amyloid (Aβ) followed by extracellular Aβ plaque deposition, and accumulation of tau with aging^10^. Therefore, identifying shared consequences of these exposures can reveal cellular drivers of LOAD. Several cellular functions have recently been implicated in LOAD including neuroinflammation, the endosomal and autophagosomal/lysosomal systems^11–18^, and altered lipid metabolism^6–9^. However, how these systems integrate with each other to result in emergence of LOAD in diverse etiological settings remains unknown.

We compared the cellular and molecular consequences of two distinct midlife risk factors for LOAD, heavy alcohol use and obesity^5,19^. A recent nationwide retrospective study found that heavy alcohol use was the strongest modifiable risk factor for dementia and AD (HR ∼ 3 and 2 respectively)^5^. Obesity during midlife is also a major risk factor for LOAD. We recently reported that early life binge alcohol increased AD pathology in adulthood^20^. This involved proinflammatory microglial activation, though the mechanism underlying neuronal amyloid accumulation was unknown. Using these two risk exposures to identify shared cellular deficits that underly LOAD pathogenesis, we found that the accumulation of neuronal lysosomal lipid (NLL) is a fundamental driver of early LOAD pathogenesis. This was caused by the loss of lysosomal acid lipase (LAL) expression, which was also found in normal aging in WT mice and healthy human brain. This phenomenon extended beyond these risk factors, with human LOAD subjects without heavy alcohol use or obesity showing widespread loss of LAL in brain, with transcriptional repression of LAL gene expression by pausing of the RNA polymerase II. This finding has significant implications for the understanding of disease progression, identification of at-risk individuals, and treatments to prevent or slow the progression of LOAD in the human population.

### LOAD midlife risk factors promote pathology by disrupting neuronal Aβ metabolism

We first assessed the impact of midlife heavy alcohol (i.e., ethanol) and diet-induced obesity on LOAD pathology in the entorhinal cortex (ENT), hippocampus, subiculum (SUB), and frontal cortex (FCX). The 3xTg-AD mouse model (APPSwe, tauP301, Psen1^tm1Mpm^) was used since it shows a progressive age-related increase in pathologic Aβ and tau species, seen first in entorhinal and hippocampal regions, followed by cortex at later ages, similar to LOAD^21–23^. Mice were treated during midlife to match the LOAD epidemiology, when Aβ pathology is ∼30-40% of peak levels in 3xTg-AD. Since mice metabolize ethanol ∼8-times more rapidly than humans^24^; a dose was used that produces average blood alcohol concentrations during intoxication similar to those in humans (average: ∼25mg/dL, peak: ∼290mg/dL)^25^. Ethanol increased pathogenic intraneuronal Aβ_1-42_ by ∼55-60% in the ENT, FCX and SUB in both sexes (Figure 1A-E). Females displayed amyloid plaques in the SUB that were increased 2-fold by ethanol (Figure 1E-F). Ethanol also increased p-tau-181, a marker for LOAD in humans^26,27^, by 40% in CA1 and FCX (Figure 1I-J). These increases occurred without any effect on expression of the human *hAPP* or *MAPT* transgenes in cortex (Extended Data Figure 1A-C). Increases in Aβ_1-42_ and p-tau-181 were accompanied by neurodegeneration, with a 2.5-fold increase in the TUNEL apoptotic stain in the ENT (Figure 1K-L) and induction of extrinsic apoptotic cell death genes implicated in AD (*TRAIL*, *caspase-8, death receptor 5* and *FADD*, Extended Data Figure 1D). WT mice were further assessed to assess the impact of AD genetics, where ethanol likewise increased total Aβ in WT ENT by 36%, (Figure 1M-N), without altering expression of mouse *APP* (Extended Data Figure 1E). Furthermore, in 3xTg-AD, ethanol had no effect on levels of cortical presenilin 1 (PSEN1) or BACE1 protein (Extended Data Figure 1F-G) and had minimal impacts on the major tau-phosphorylating kinases (pSer9-GSK3β, pTyr216-GSK3β, p-PKA) and the tau phosphatase PP2A (Figure 2H-L). Midlife likewise obesity increased Aβ_1-42_ at 11 months 0.5 to 2.5-fold in ENT, FCX, and SUB (2.4-fold) (Figure 1O-T). Similar to ethanol, this occurred without any differences in expression of the *hAPP* transgene (Extended Data Figure 1N-O) or levels of PSEN1 or BACE1 (Extended Data Figure 1P-Q). Obese mice also had increased p-tau181 in CA1 (41%, *p=*0.05, Figure 1U-V) with no changes in expression of the *hMAPT* transgene (Extended Data Figure 1R), nor tau phosphorylating enzymes GSK3β, p-PKA, CDK5, p16 (Extended Data Figure 1S-Z).

**Figure 1.**
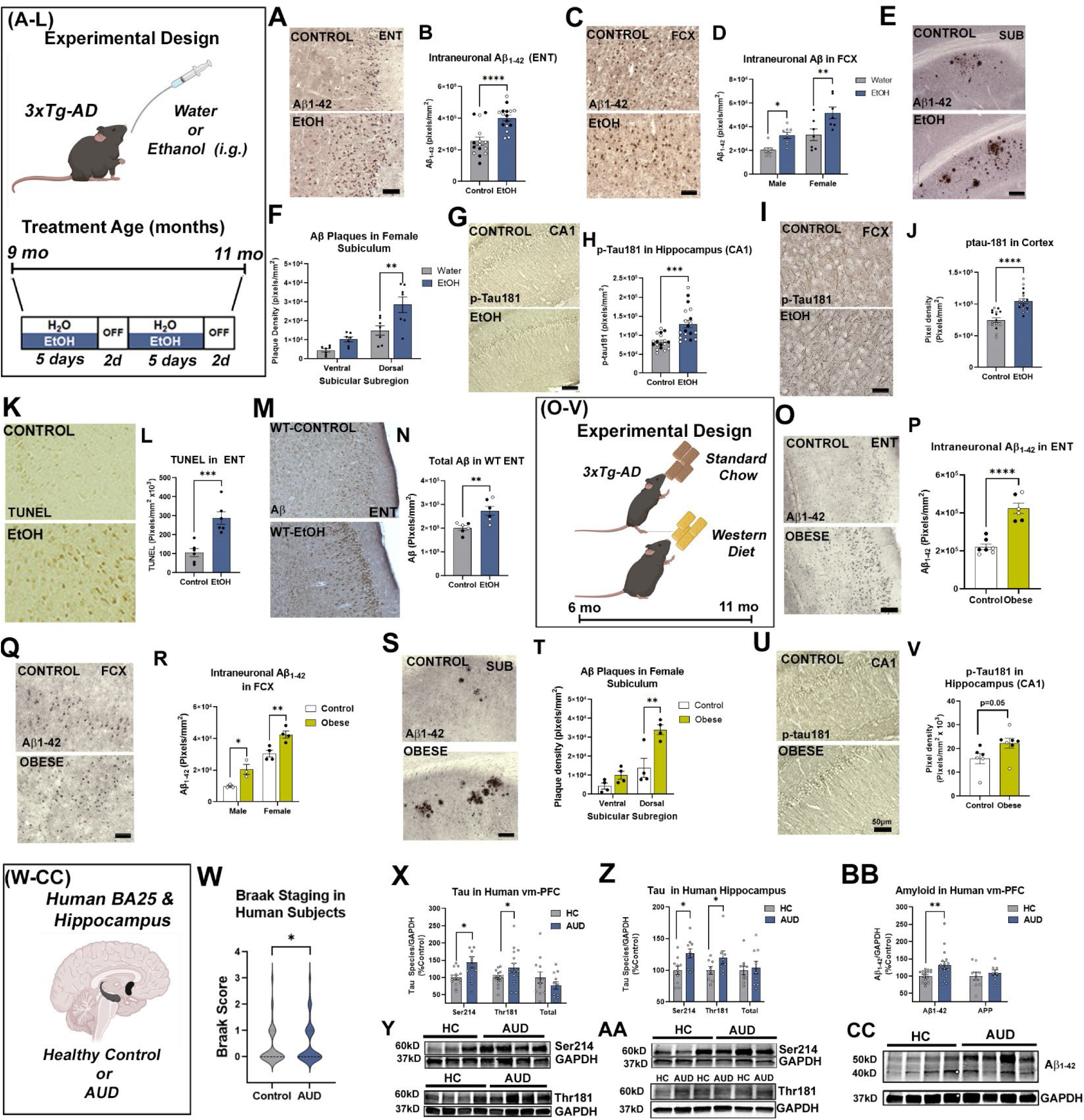
Midlife alcohol and obesity increase LOAD pathology. **(A, B)** Ethanol increased intraneuronal Aβ_1-42_ in ENT 57%. N=15 control, 16 ethanol. **(C, D)** Ethanol increased Aβ_1-42_ in FCX in both sexes. F_1,26_=16.83, p=0.0004, \**p*<0.05, \*\**p*<0.01 Sidak’s post-test. Scale: 100µm **(E, F)** Ethanol increased Aβ plaques in the female subiculum by 91% (dorsal) and 127% (ventral). Scale bar: 200µm. Ethanol **i**ncreased p-tau181 in **(G)** CA1 and **(H)** FCX. N=18 control, 19 ethanol. **(I, J)** 40% increase in p-tau181 in FCX. Scale bar: 50µm. **(K, L)** Ethanol increased TUNEL+ neurodegeneration in ENT by 3-fold. N=6/group. **(M, N)** 5 weeks of ethanol increased total Aβ in ENT of WT 10-mo mice. **(O, P)** Obesity increased ENT Aβ_1-42_ by 47%. N=7/group. **(Q, R)** Obesity increased intraneuronal Aβ_1-42_ in FCX 2-fold in males and 40% in females. Sex: F_1,10_=89.2, p<0.0001, Obesity: F_1,10_=25.2, p=0.0005 **(S, T)** 2.4-fold increase in Aβ plaques in the dorsal subiculum of obese females. Scale bar: 200µm. \*\**p*<0.01, Sidak’s multiple comparison test. **(U, V)** 41% increase in p-tau181 in CA1 in obese mice. N=6 Control, 7 obese. *p*=0.05. **(W)** Braak staging scores were slightly increased in AUD. \**p*<0.05, paired *t-*test. **(X, Y)** Human AUD vm-PFC had a ∼50% increase in p-tau214 and a ∼30% increase in p-tau181 with no change in total tau. \*\**p*<0.01, paired *t-*test, N=16/group. **(Z, AA)** Human AUD hippocampus showed a 27% increase in p-tau214 and a 20% increase in p-tau181 by Western blot. \*\**p*<0.01. **(BB, CC)** Aβ_1-42_ was increased in AUD vm-PFC by 31%. N=17/group. *t*-test. \**p*<0.5 \*\**p*<0.01 \*\*\**p*<0.001. Males-open circles, Females-closed circles.

**Figure 2.**
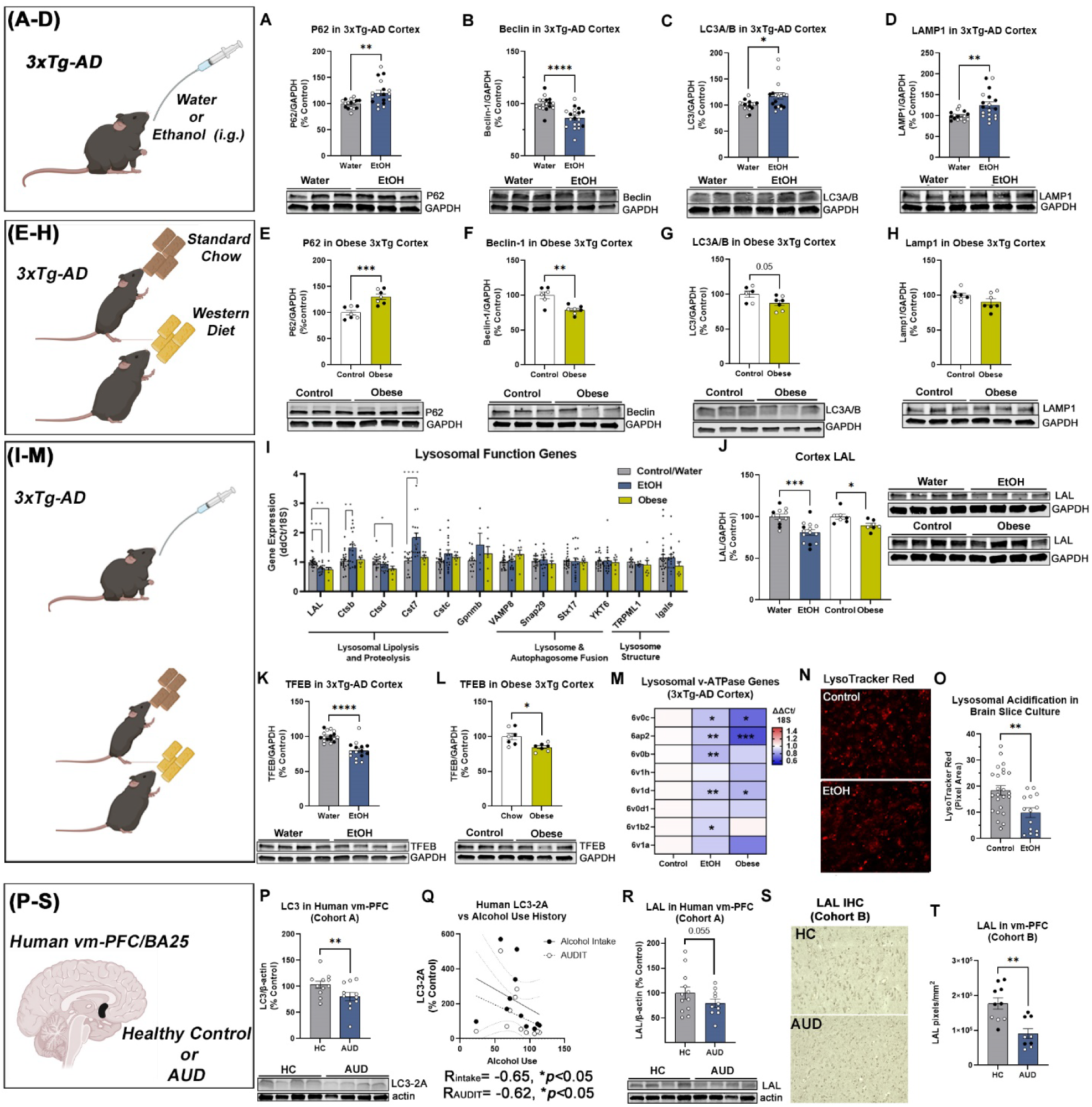
Ethanol and obesity disrupt autophagic flux and lysosomal function in 3xTg-AD mice. Ethanol **(A)** increased P62 by 20%, **(B)** reduced Beclin-1 by 14%, **(C)** increased LC3 by 20%, and **(D)** increased LAMP1 by 26% in 3xTg-AD cortex. N=15 control, 18 ethanol. \**p*<0.05, \*\**p*<0.01, \*\*\*\**p*<0.0001, *t-*tests. Males-open circles, females-closed circles. Obesity **(E)** increased P62 by 33% **(F)** reduced Beclin 21%, and **(G)** reduced LC3 by 12.5% in 3xTg-AD cortex. **(H)** No change in LAMP1 was seen in obese 3xTg-AD cortex. N=6 control, 7 diet-induced obesity. \*\**p*<0.01, \*\*\**p*<0.001, *t-*tests. **(I)** LAL (*LIPA*) gene expression was reduced by both EtOH and obesity. **(J)** Reductions in LAL protein caused by ethanol (19%) and obesity (10%). Both ethanol **(K)** and obesity **(L)** reduced in TFEB protein in 3xTg-AD cortex (ethanol-20%, obesity-15.4%). \**p*<0.05, *t-*test. **(M)** Reduced expression of lysosome acidifying v-ATPases genes by both ethanol and obesity. **(N)** Lysosomal acidification in HEBSC slice cultures was reduced by ethanol (100mM, 4 days) by **(O)** 50% as measured by LysoTracker red. **(P-S)** Altered autophagy and reduced LAL in post-human AUD vm-PFC/BA25 in two cohorts. **(P)** LC3-II was reduced by ∼20% in AUD vm-PFC and **(Q)** was correlated negatively with lifetime alcohol use and AUDIT score. **(R)** In human cohort A, a reduction in LAL was found in AUD by Western blot that approach statistical significance. *p*=0.055. **(S, T)** IHC on subjects from cohort B found a 50% reduction in LAL in human AUD vm-PFC. \*\**p*<0.01, paired *t-*test.

Findings in rodents were consistent with human postmortem ventromedial prefrontal cortex (vm-PFC/BA25) and hippocampus from individuals with alcohol use disorder (AUD) or moderate-drinking age-matched controls. Though none of these individuals had a co-morbid diagnosis of AD, AUD brains had higher Braak scores than controls (Figure 1W) and increased levels of p-tau214 and p-tau181 in vm-PFC (Figure 1X-Y, ∼50% and 30%, respectively) and hippocampus (Figure 1Z-AA, 20-27%) without increases in total tau. Aβ_1-42_ was also increased in AUD vm-PFC (31%, Figure 1BB-CC), with no changes in levels of amyloid precursor protein (APP). Similar to findings in 3xTg-AD, gene expression of *APP*, *MAPT*, *GSK3* and *CDK5* were not changed in AUD hippocampus (Extended Data Figure 1AA). Tau-phosphorylating isoforms of GSK3 were increased in AUD BA25 (Extended Data Figure 1BB-CC), while levels of Aβ_1-42_ producing BACE1 protein were unchanged (Extended Data Figure 1DD). Together, this supports that both midlife alcohol and obesity promote LOAD pathology without altering Aβ or tau gene expression or levels of Aβ modifying enzymes.

**Extended Data Figure 1.**
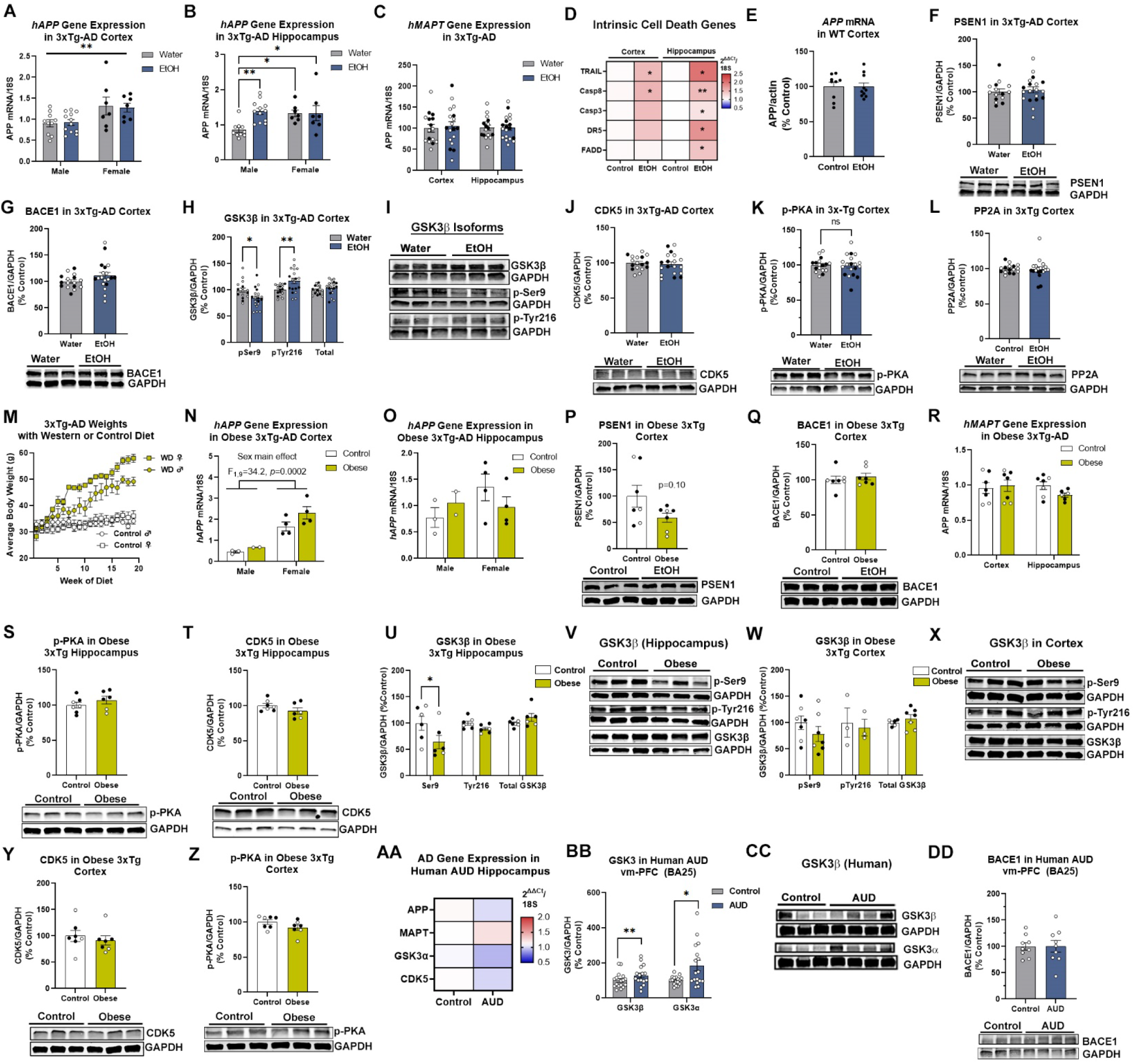
Neither chronic ethanol nor obesity had a significant impact on transgene expression, Aβ or Tau processing machinery. **(A)** Ethanol had no effect on expression of the human *APP* (*hAPP*) transgene in cortex of 3xTg-AD mice. **(B)** Ethanol increased *hAPP* in male 3xTg-AD hippocampus by 38%, with no change in females. Ethanol x sex, F_1,32_=10.6, *p=*0.03. **(C)** No effect of ethanol on *hMAPT* in cortex or hippocampus. **(D)** Ethanol increased intrinsic cell death genes in 3xTg-AD cortex and hippocampus. **(E)** Ethanol had no impact on *APP* in WT mice. **(F)** Ethanol had no effect on PSEN1 in 3xTg-AD mice. **(G)** Ethanol had no effect on BACE1 levels in 3xTg-AD cortex. **(H, I)** Ethanol reduced the inactive pSer9 isoform by 15.6% and increased active p-Tyr216 by 17% without significantly total GSK3β in 3xTg-AD mice. No increases in cortical levels of **(J)** CDK5, nor **(K)** p-PKA were found. **(L)** No effect of ethanol on PP2A in 3xTg-AD cortex. *p=*0.28. **(M)** Weights of 3xTg-AD mice receiving Western diet across midlife. **(N)** A main effect of sex on *hAPP* F_1,9_=34.2, *p=*0.0002, with no effect of obesity. **(O)** No change in *hAPP* expression in 3xTg-AD hippocampus. **(P)** No significant differences in PSEN1 or **(Q)** BACE1 in obese mice. **(R)** No change in *hMAPT* with obesity. **(S)** No changes in p-PKA or **(T)** CDK5 in hippocampus. **(U, V)** Total and active GSK3β were unchanged in obese hippocampus, with a ∼50% reduction of inactive p-Ser9 GSK3β. Obesity did not change cortical **(W, X)** GSK3β **(Y)** CDK5 or **(Z)** p-PKA. N=7/group. **(AA)** Expression of *APP, MAPT, GSK3,* and *CDK5* were unchanged in human AUD hippocampus. **(BB-CC)** Increased GSK3β and GSK3α were in human AUD vm-PFC. AUD Cohorts 1 and 2, N = 17 age-matched healthy control and AUD subjects. **(DD)** No change in BACE1 protein in human AUD vm-PFC. AUD Cohort A, N= 9 age-matched healthy control and AUD subjects. \**p*<0.05, \*\**p*<0.01 paired *t-*tests.

### Disruption of autophagy and lysosomal function by heavy alcohol and obesity during midlife

Since neurochemical findings suggested midlife LOAD risk exposures disrupt neuronal Aβ metabolism, we assessed the autophago-lysosomal system, the main site of intraneuronal Aβ degradation^28,29^. Midlife ethanol increased p62 (20%), reduced the autophagy initiator Beclin (20%), increased the autophagosome elongation factor LC3-II (20%), and increased the mature lysosome protein LAMP1 (26%) and increased phosphorylated mTOR (p-mTOR, 2-fold, Ser4228) (Figure 2A-D, Extended Data Figure 2A, 2-fold) in 3xTg-AD cortex. In WT mice ethanol increased P62 in females, reduced Beclin in both sexes, and reduced LC3 in both sexes, with LAMP1 accumulation in females (Extended Data Figure 2E-H). These findings are consistent with a loss of autophagic flux secondary to lysosomal dysfunction and a secondary reduction in autophagy initiation (Extended Data Figure 2B). Diet-induced obesity also disrupted autophagic flux, though differently than ethanol (Extended Data Figure 2C). Obesity increased p62 (33%), reduced Beclin (21%), and reduced LC3 (12.5%), total without changing LAMP1 (Figure 2E-H) or p-mTOR Ser4228 (Extended Data Figure 2D). In 11-month obese WT mice, no significant changes in P62, Beclin, LC3 or LAMP1 were seen (Extended Data Figure 2I-L).

To further assess lysosomal function, we measured the expression of lysosomal lipases and proteases, regulators of lysosome/autophagosome fusion, and lysosomal structural genes (Figure 2I). Neither ethanol nor obesity significantly altered the expression of facilitators of lysosome/autophagosomes fusion nor regulators of lysosomal structure. However, both LOAD risk exposures reduced expression of the LAL gene (*LIPA*, 20-25%) and protein (10-20%, Figure 2J). This occurred with reductions in TFEB, a regulator of lysosomal gene expression^30^ (Figure 2K-L) and reduced lysosome acidifying v-ATPases (Figure 2M). Similar reductions in TFEB and lysosomal v-ATPases were seen in WT female mice with ethanol (Extended Data Figure 2M-N). Of note, TFEB was expressed robustly in cortical neurons (Extended Data Figure 2O). We then found ethanol reduced lysosomal acidification in the ENT of hippocampal-entorhinal brain slice cultures (HEBSC, Figure 2N-O). In human AUD brain, LC3 was reduced 20% and correlated negatively with lifetime alcohol use and AUDIT score (Figure 2Q). LAL protein levels were also reduced in AUD vm-PFC (Figure 2R-T). Other AD-associated regions also showed reduced LAL in AUD including ENT (39%), CA1 (27%), and SUB (54%) (Extended Data Figure 2P-Q). Together this indicates these two common midlife LOAD risk factors disrupt autophago-lysosomal flux and reduce LAL in rodents and humans.

**Extended Data Figure 2.**
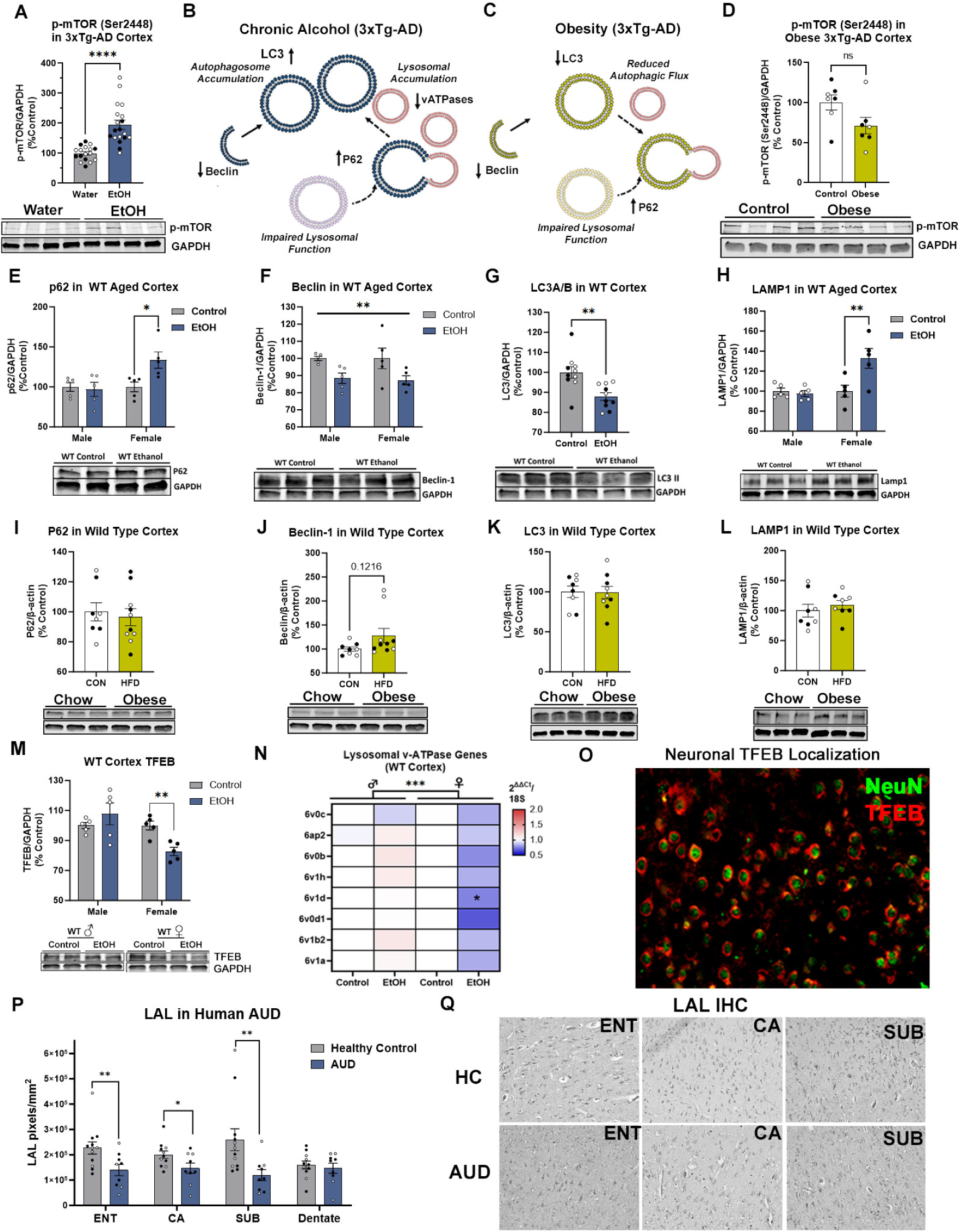
Midlife alcohol and obesity alter autophagy and LAL in rodents and humans. **(A)** Ethanol increased p-mTOR (Ser2448) by ∼2-fold in 3xTg-AD cortex. **(B)** Schematic summarizing reduced autophagic flux in 3xTg-AD mice by ethanol. **(C)** Schematic illustrating disruption of autophagic flux caused by midlife obesity. **(D)** Trend toward reduced p-mTOR (Ser2448) in obese 3x-Tg-AD cortex. **(E-H)** Autophagy markers in WT cortex of 10-mo mice treated with ethanol for 5 weeks. **(E)** Ethanol increased P62 protein levels in females by 33%, \**p=*0.018, and did not alter levels in males. **(F)** Main effect of ethanol on Beclin-1 protein. F_1,16_=11.00, \*\**p*=0.004. There was a significant decrease in Beclin-1 protein in males (12% reduction, \*\**p*=0.008) and a trend toward a reduction in females (13%, p=0.08, *t-*test). **(G)** Ethanol reduced LC3A/B by 12%, \*\**p*=0.0026, and **(H)** increased LAMP1 levels in females (∼33%, \*\**p*=0.0035). Obesity had no effect on **(I)** P62 levels, **(J)** Beclin, **(K)** LC3, or **(L)** LAMP1. **(M)** Ethanol caused a loss of TFEB in female WT cortex. **(N)** Reduction of lysosomal v-ATPase gene expression in the cortex of WT female mice treated with ethanol. \**p*<0.05*, **p*<0.01, \*\*\**p*<0.001. **(O)** Co-immunofluorescent labeling of TFEB and neuronal marker NeuN showing robust localization of TFEB in neurons. LAL was reduced in AUD (Cohort B) compared to age-matched controls as measured by IHC in **(P, Q)** ENT-39%, CA1-27%, and subiculum-54% with no change was seen in the dentate. \**p*<0.05, \*\**p*<0.01 Paired *t-*tests.

### Enhancement of lysosomal lipid accumulation within Aβ+ neurons with obesity and ethanol

We next determined if lysosomal lipid is associated with accumulation of Aβ. Cytosolic lipid was increased with both LOAD risk factors in FCX (Figure 3A-B, obesity: 80%, ethanol: 38%) with greater increases were in the ENT, which develops amyloid pathology prior to the FCX (obese: 120%, ethanol: 88%). Notably, significant lipid increases were found in lysosomes in FCX (Figure 3C, obesity: 64%, ethanol: 45%) and ENT (obesity: 1.8-fold, ethanol: 2.3-fold). Lysosomal lipid was increased in Aβ+ neurons by obesity and ethanol (Figure 3D, FCX ∼75%, ENT-obese: 3.8-fold, ethanol: 3.3-fold). Of note, lipid accumulation was also found in microglia consistent with recent reports (Extended Data Figure 3A-C)^31^. Lysosomal lipid was also associated with an increase in lysosome number (Figure 3E). As above, ethanol and obesity increased Aβ cortex (Figure 3F, FCX-∼55%, ENT-obese: 2.1-fold, ethanol 59%). Of note, these increases in Aβ were in the extra-lysosomal cytosol (Figure 3G-H) with reductions in the Aβ lyso:cyto ratios (Figure 3I). With each exposure, intraneuronal Aβ was strongly correlated with neuronal lysosomal lipid (NLL, Figure 3K-L, FCX-obese: R^2^=0.72, \*\*\**p=*0.0004, ethanol R^2^=0.63, \*\**p=*0.003; ENT-obese: R^2^=0.83, \*\*\**p=*0.0005, ethanol: R^2^=0.81, \*\*\*\**p*<0.0001). WT mice treated with ethanol likewise had increased total (83%, Extended Data Figure 3D-E) and lysosomal lipid (2.5-fold, Extended Data Figure 3F) with a strong correlation between total Aβ and total lipid (Extended Data Figure 3G, R=0.9, \*\*\*\**p*<0.0001), indicating AD genetics are not required. CA1 p-tau was increased by ethanol, with no change in the lyso:cyto ratio suggesting an alternate mechanism (Extended Data Figure 3H-J). We did not find evidence that ER stress, which promotes lipid droplet formation, was involved (Extended Data Figure 3K). Neither did we find robust changes in lipid metabolism genes or cytosolic lipases with ethanol, obesity, or aging that could explain the enhanced lysosomal lipid profile in both sexes (Extended Data Figure 3L-O). However, both exposures altered the expression of lipid-droplet coating perilipins (PLINs) and reduced expression of lipid efflux transporters *ABCG1* and *ABCA1* (Extended Data Figure 3L).

**Figure 3.**
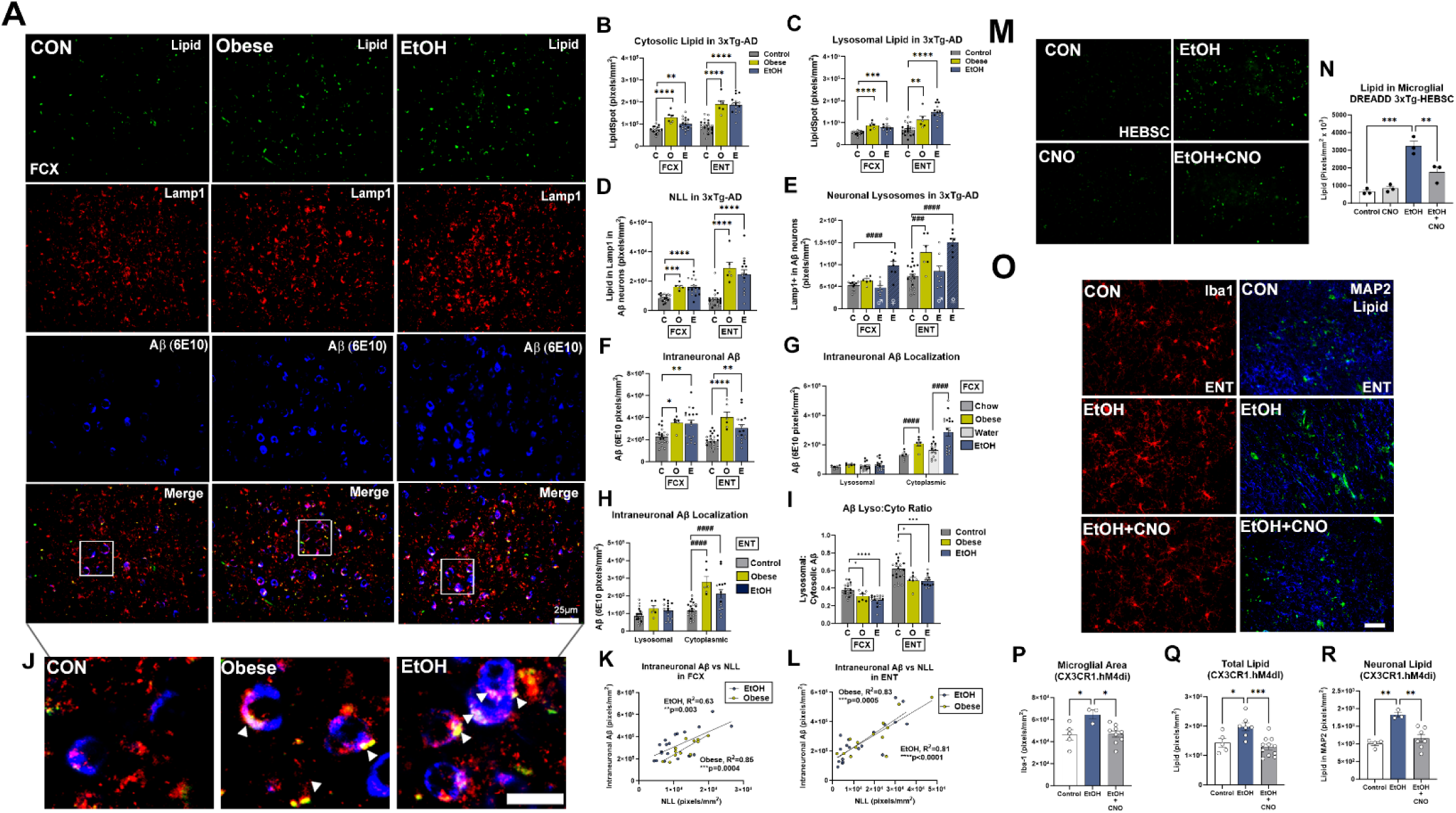
Increased lysosomal lipid within Aβ+ neurons with obesity and ethanol that is enhanced by proinflammatory microglia. **(A)** Triple immunofluorescence (IF) for cytosolic lipid (LipidSpot, green), LAMP1 (red), and Aβ (blue). Normal chow and water gavage groups were combined when no differences were present. Scale bar: 25µm. **(B)** Cytosolic lipid was increased by obesity (FCX: 80%, ENT: 120%) and EtOH (FCX: 38%, ENT: 80%). EtOH effect F_2,40_=20.07, p<0.0001. Obesity: F_2,41_=35.84, p<0.0001. **(C)** Increased lipid within lysosomes in FCX (obesity: 64%, ethanol: 45%, F_2,29_=16.09, p<0.0001, treatment effect) and ENT (obese: 1.8-fold, ethanol: 2.3-fold, F_2,41_=40.2, p<0.0001, treatment effect). **(D)** Increased lysosomal lipid in Aβ+ neurons in FCX (obese: 79%, ethanol: 75%, F_2,41_=15.73, p=0.0001, treatment effect) and ENT (obese: 3.8-fold, ethanol 3.3-fold, F2,40=32.1, p<0.0001, treatment effect). **(E)** Increased lysosomes in Aβ+ neurons with ethanol (2-fold FCX and ENT) and obesity (ENT: 76%). F_3,32_=16.46, p<0.0001, treatment**. (F)** Increased intraneuronal Aβ in FCX (obesity: 56%, ethanol: 52%, F_2,41_=7.5, p=0.002) and ENT (obesity: 2.1-fold, ethanol: 59%, F_2,41_=15.3, *p=*0.0001). Extra-lysosomal cytoplasmic levels of Aβ were increased in **(G)** FCX in (obese: 61%, F_1,20_=19.52, *p=*0.0003; ethanol: 1.8-fold, F_1,59_=14.38, *p=*0.0004) and **(H)** ENT (obese: 2.2-fold, ethanol: 1.9-fold, F_2,82_=20.38, *p=*0.00001). **(I)** Reduction of lysosomal:extra-lysosomal cytoplasmic (lyso:cyto) Aβ ratios. FCX-F_2,41_=14.50, *p*<0.0001, ENT-F_2,40_=8.788, *p*=0.0007 **(J)** High magnification image showing increased NLL in Aβ+ neurons. Scale: 10µm. Strong positive correlations of NLL with intraneuronal Aβ in **(K)** FCX and **(L)** ENT with obesity and ethanol. **(M, N)** Ethanol-induced increases in cytosolic lipid in HEBSCs was prevented by microglial inhibition (hM4di). **(O, P)** Chemogenetic inhibition of proinflammatory microglia (CX3CR1Cre^ERT2^.hM4di) prevented ethanol-induced **(P)** microglial activation, **(Q)** increases in total cytosolic lipid and **(R)** NLL accumulation in the ENT. \**p*<0.05, \*\**p*<0.01, \*\*\**p*<0.001, \*\*\*\**p*<0.0001, Dunnett’s post-test. #*p*<0.05, ##*p*<0.01, ###*p*<0.001, ####*p*<0.0001, Sidak’s post-test. Open circles: males, filled circles: females.

Neither ethanol nor obesity caused convergent expression changes of common upstream regulators of lipid metabolism or production (Extended Data Figure 3M-O). However, we and others have reported that proinflammatory microglia activation is associated with intraneuronal Aβ accumulation^20,32^. Therefore, we assessed if proinflammatory microglia promote NLL. HEBSCs from 3xTg-AD mice were transfected with a Gi inhibitory DREADD (AAV9.CD68.hM4di) to inhibit proinflammatory signaling as we previously reported^33,34^. Inhibition of microglial proinflammatory blocked ethanol-induced increases in cytosolic lipid (Figure 3M-N). This also occurred *in vivo* using CX3CR1Cre^ERT2^.hM4di mice. Ethanol promoted proinflammatory activation of microglia as well as increases in total and neuronal lipid that were blocked by microglial inhibition (Figure 3O-R). Together, this suggested NLL promotes intraneuronal Aβ pathology that is enhanced by proinflammatory microglia. Therefore, we next determined if loss of LAL causes NLL to drive Aβ accumulation.

**Extended Data Figure 3.**
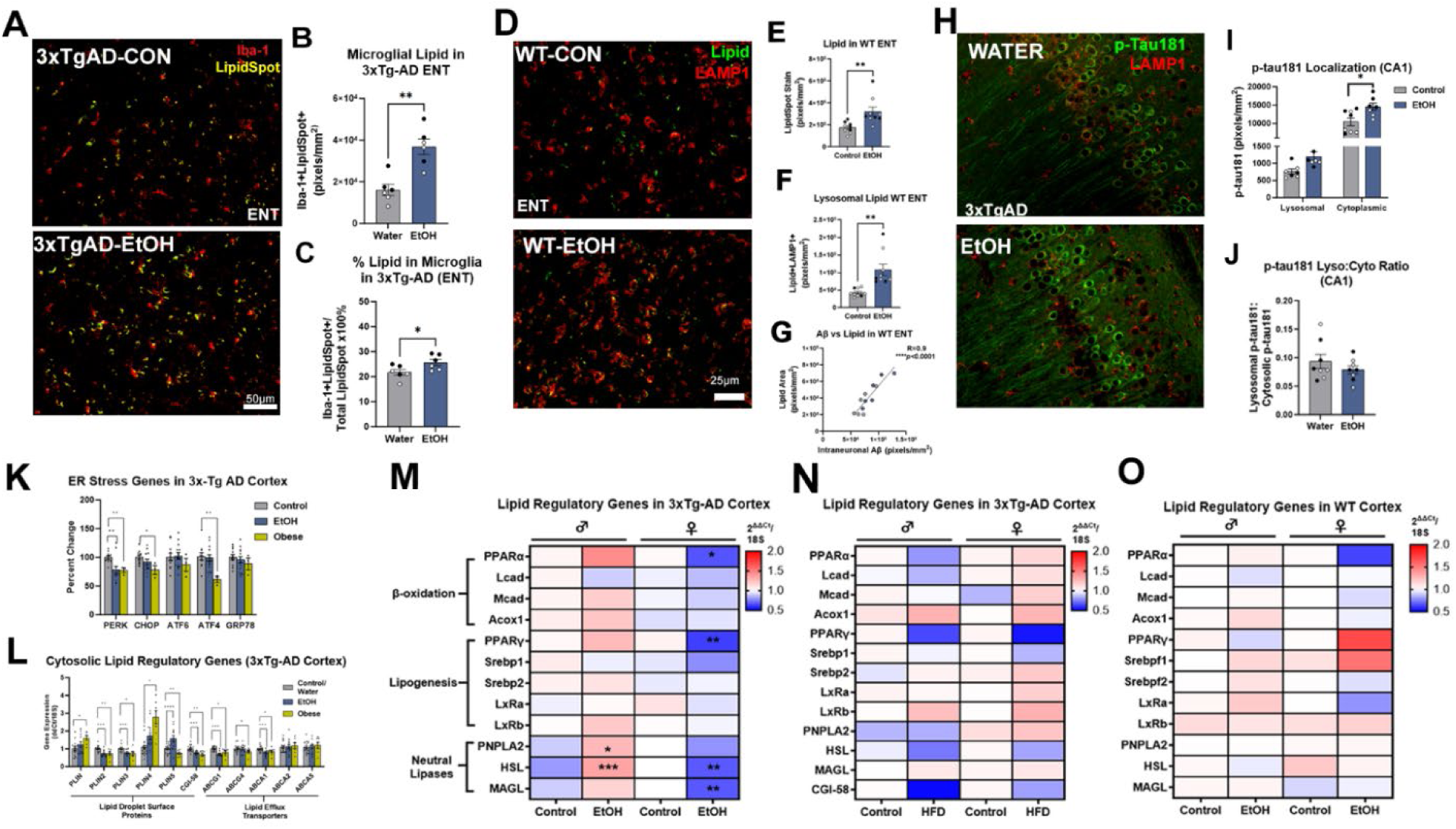
Lipid metabolic changes with ethanol or obesity in 3xTg-AD and WT mice. **(A)** Representative image of microglial lipid levels in 3xTg-AD mice after chronic ethanol. Red: Iba-1, yellow-LipidSpot stain. **(B)** Quantification of lipid within Iba-1+ found an ∼2-fold increase in microglial lipid after ethanol treatment. **(C)** The percentage of total lipid localized to microglia was slightly increased after ethanol. \**p*<0.05. **(D)** Representative image of lysosomal (LAMP1) lipid staining in WT mice treated with water or ethanol. Scale bar: 25µm. **(E)** Quantification found ethanol increased total lipid in WT mice. **(F)** Ethanol increased the level of lysosomal lipid by ∼2-fold. \*\**p*<0.01. **(G)** A strong correlation was found between the level of lipid and Aβ in ENT of WT mice. R=0.9. **(H)** Co-IF for p-tau 181 and LAMP1 in 3xTg-AD CA1 after ethanol. Scale bar: 25µm. **(I)** primarily cytosolic p-tau-181 was increased by ethanol lysosomal p-tau 181 was increased after ethanol. **(J)** The p-tau 181 lyso:cyto ratio was not significantly reduced by ethanol. **(K)** Neither ethanol nor obesity caused an upregulation in ER stress related genes. **(L)** Gene expression of lipid droplet associated perilipins (PLINs), CGI-58, and lipid efflux transporters showed alterations in PLIN expression and reductions in ABCG1 and ABCA1 by both ethanol and obesity. Expression of major regulators of lipid metabolism and cytosolic lipases showed no robust pattern shared by both males and females after **(M)** ethanol or **(N)** diet-induced obesity in 3xTg-AD mice. **(O)** Ethanol had no robust effect on expression of key lipid metabolism genes or cytosolic lipases in WT mice. \**p*<0.05*, **p*<0.01, \*\*\**p*<0.001, \*\*\*\**p*<0.0001.

### LAL is lost with aging and promotes NLL accumulation to drive Aβ accumulation

LOAD pathology emerges in humans and 3xTg-AD mice with aging, suggesting the loss of underlying resilience mechanisms. To determine if NLL accumulation is an age-related risk factor for LOAD, we assessed WT mice across aging. At 3 months, very little intracellular lipid was seen (Figure 4A). However, by 20 months profound increases lipid were found in neurons and lysosomes (Figure 4B-D) with some sex differences. In FCX, cytosolic lipid increased 280-fold from 3 to 20 months in females, with males reaching a stable 19-fold increase by 12 months (Extended Data Figure 4A). Neuronal and lysosomal lipid also increased greatly in male and female FCX (Figure 4E-F) with similar changes seen in ENT for total lipid (Extended Data Figure 4B), neuronal lipid, and lysosomal lipid (Figure 4G-H). No differences were seen in total or neuronal LAMP1 (Extended Data Figure 4C-D). However, robust increases in NLL were found, that reached higher levels in females (Figure 4I-J). Notably, expression of *PLIN3* and *PLIN4* increased in both sexes (Figure 4K-L) as did the lipogenic factor *Srebp2* (Figure 4M). In general, females, which show increased for LOAD, had a more active lipid regulatory profile than males (Extended Data Figure 4I-J). This includes increases in *LXRα* and metabolic genes *LCAD and MCAD* (Extended Data Figure 4I-J). In ENT, total LAL and neuronal LAL declined ∼20-30% with age (Figures 4N-O and S4E), while in FCX, a main effect of age on LAL was found with females having lower levels at 3 months than males (Extended Data Figure 4F, F_2,29_=20.64, *p*<0.001). Age-related reductions in neuronal LAL in FCX were less robust than in ENT, though males showed a slight decline from 3 to 12 months (Extended Data Figure 4G). At 12 and 20 months, NLL was negatively correlated with neuronal LAL (Extended Data Figure 4H, R= -0.38, \**p*<0.05). Total Aβ in WT ENT appeared to lag slightly behind LAL loss, as it increased from 3 to 20 months (25%, Figure 4P-Q), was positively correlated with NLL (Figure 4R, R= 0.45, \*\**p*<0.01), and negatively correlated with neuronal LAL (Figure 4S, R= -0.33, \**p*<0.02). Together, this indicates that in normal aging, NLL increases and is strongly associated with Aβ accumulation, similar to findings in AD mice.

**Figure 4.**
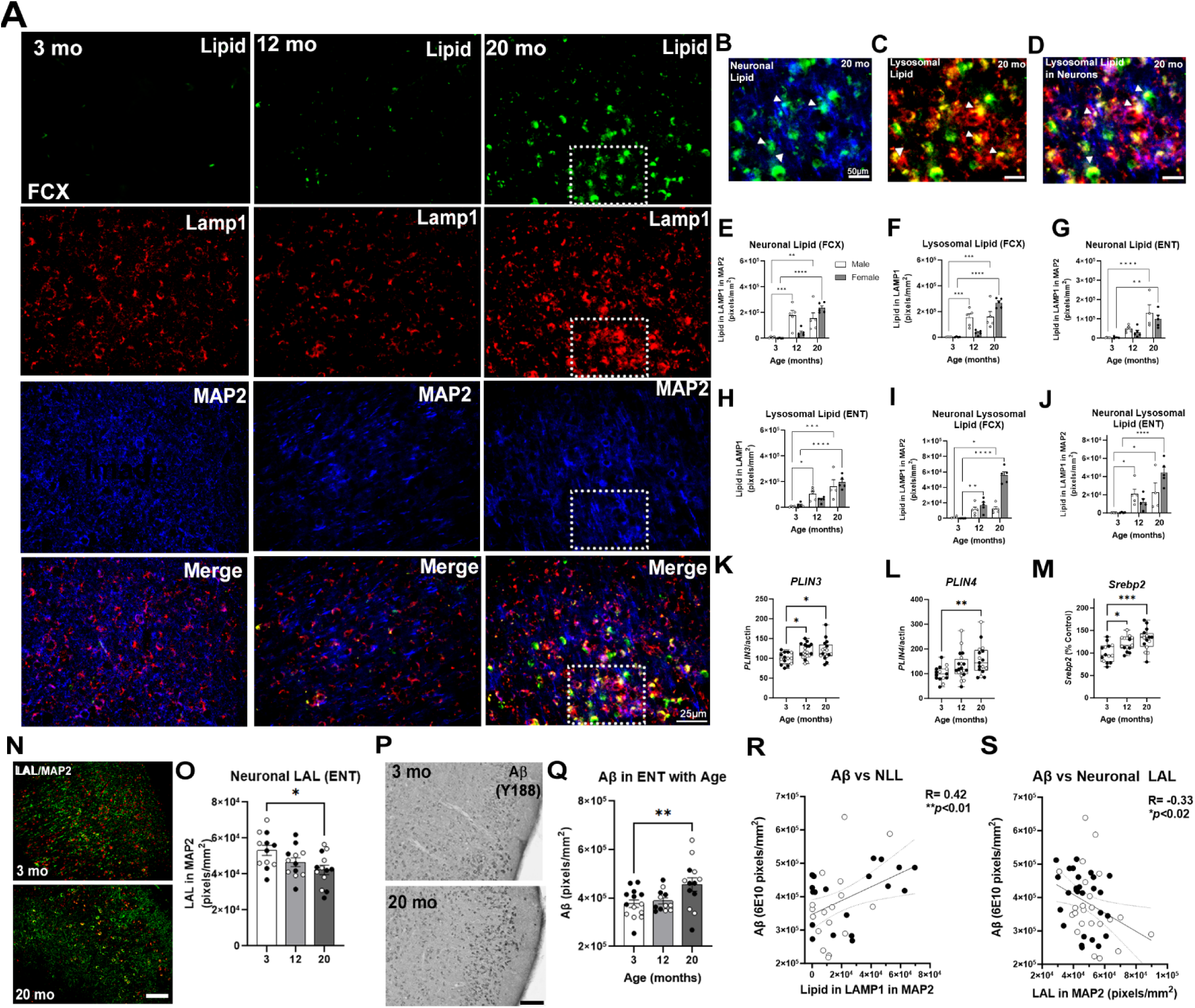
Loss of neuronal LAL and accumulation of neuronal lysosomal lipid with aging in wild-type mice. **(A)** Representative images of cytosolic lipid (LipidSpot), lysosomes (LAMP1), and neurons (MAP2) at 3, 12, and 20 months of age in frontal cortex (FCX). Higher magnification images of **(B)** Neuronal lipid, **(C)** lysosomal lipid, and **(D)** neuronal lysosomal lipid at 20 months. Profound aging-related increases in FCX **(E)** neuronal lipid, **(F)** lysosomal lipid, and **(G)** NLL, as well as ENT **(H)** neuronal lipid, **(I)** lysosomal lipid, **(J)** and NLL from 3 to 20 months. 2-way ANOVAs, \**p*<0.05, \*\**p*<0.01, \*\*\**p*<0.001, \*\*\*\**p*<0.0001, Dunnett’s post-test. Increased expression of **(K)** *PLIN3,* 1-way ANOVA F_2,48_=4.85, *p*<0.02, and **(L)** *PLIN4,* F_2,49_=4.85, *p*<0.02 and **(M)** *Srepb2* expression increases with age. F_2,42_=8.45, *p*<0.001. \**p*<0.05, \*\**p*<0.01, \*\*\**p*<0.001, Sidak’s post-test. **(N-O)** Images of total and neuronal (within MAP2/green) LAL (red) with a 30% loss of neuronal LAL in ENT with age. **(P-Q)** In WT mice, a 30% increase in total Aβ (6E10) was found in ENT from 3 to 20 months. **(R)** Total ENT Aβ was positively correlated with neuronal lysosomal lipid in WT ENT **(S)** Negative correlation between Aβ and neuronal LAL. Open circles: males, filled circles: females.

**Extended Data Figure 4.**
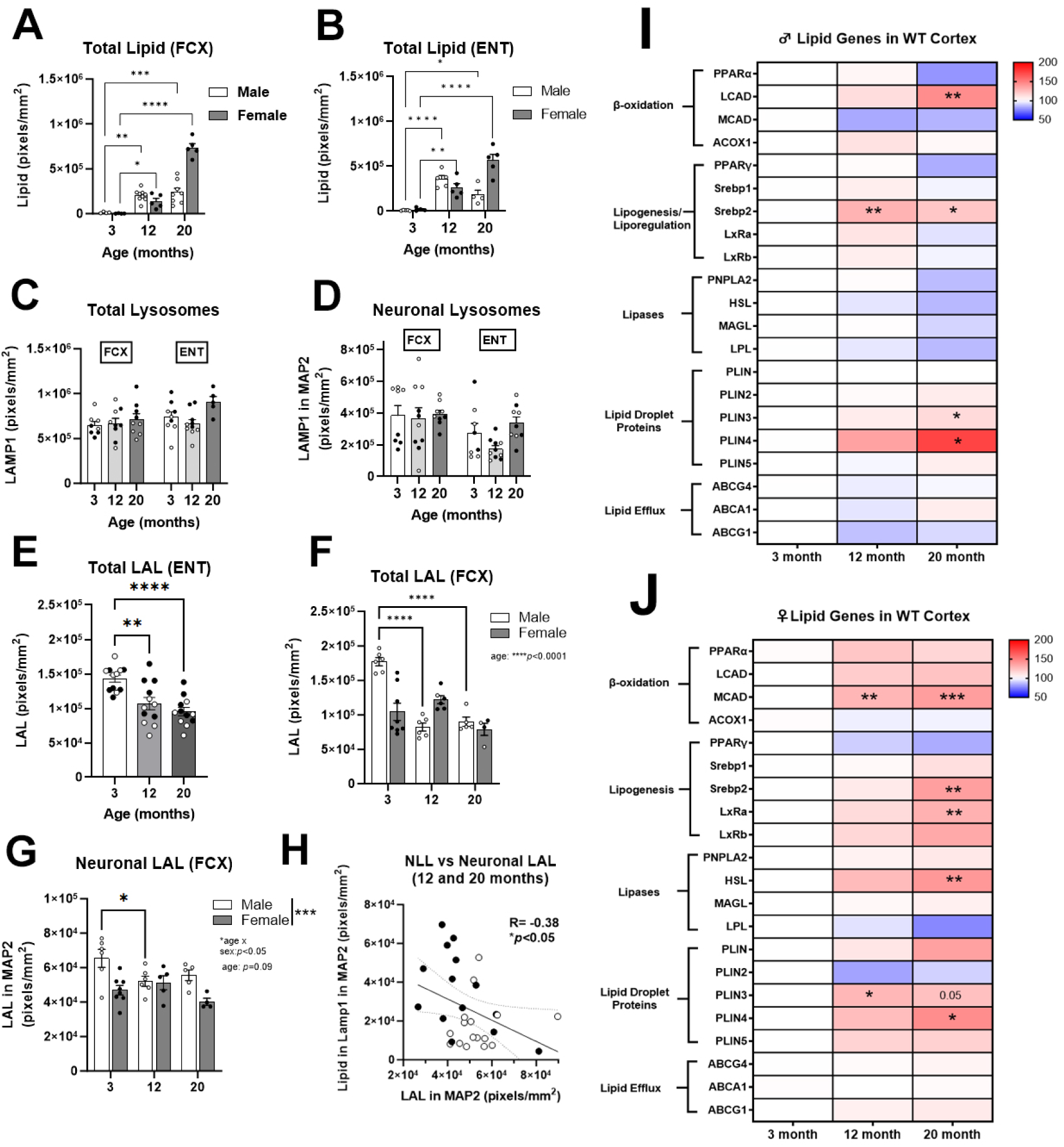
Age related changes in neuronal lipids and metabolic enzymes. Total cytosolic lipid increased dramatically in **(A)** FCX. F_2,29_=84.71, *p*<0.001 and **(B)** ENT. No changes in **(C)** total or **(D)** neuronal lysosomes were found in FCX or ENT with age. **(E)** total LAL in ENT declined ∼30% from 3 to 20 months. In FCX, a reduction in **(F)** total LAL was found, with **(G)** sex-specific reductions in neuronal LAL. **(H)** Across all subjects a negative correlation was found between neuronal lysosomal lipid and neuronal LAL. R= -0.36, \**p*<0.05. Expression of several lipid-regulatory genes in WT **(I)** male and **(J)** female cortex with age.

We next investigated the direct role of LAL loss on Aβ pathology. Assessment of the spatial-temporal relationship between LAL and Aβ in AD mice found that neuronal LAL declined with age and preceded accumulation of Aβ AD mouse cortex (Figure 5A-B). At 11 months, regions with the highest levels of Aβ pathology (e.g., SUB) had the lowest levels of LAL (Figure 5C-D). Both midlife ethanol and obesity reduced total and neuronal LAL (Figure 5E-H) with strong negative correlations between Aβ and LAL across all regions assessed (Figure 5I-J). Inhibition of LAL activity with LAListat (LALi) in 3xTg-AD HEBSC caused a concentration-dependent increase in cytosolic lipid (Figure 5K-L) and intraneuronal Aβ (Figure 5M-N) in the ENT. Induction of master lysosomal transcription factor TFEB with genistein (Extended Data Figure 5A-B) and GLP-1/GIP signaling with DA4-JC (Extended Data Figure 5C-D) did not abolish ethanol-induced increases in cytosolic lipid. However, recombinant LAL/sebelipase (rLAL) blocked the 2-fold increase in cytosolic lipid caused by ethanol (Figure 5O-P), reduced baseline Aβ levels by 50% and abolished the 2-fold increase in Aβ caused by ethanol (Figure 5Q-R), indicating LAL loss causes Aβ accumulation. Thus, LAL loss precedes Aβ accumulation with age, LAL levels are negatively correlated with Aβ *in vivo,* and LAL inhibition directly enhances Aβ levels *ex-vivo*.

**Figure 5.**
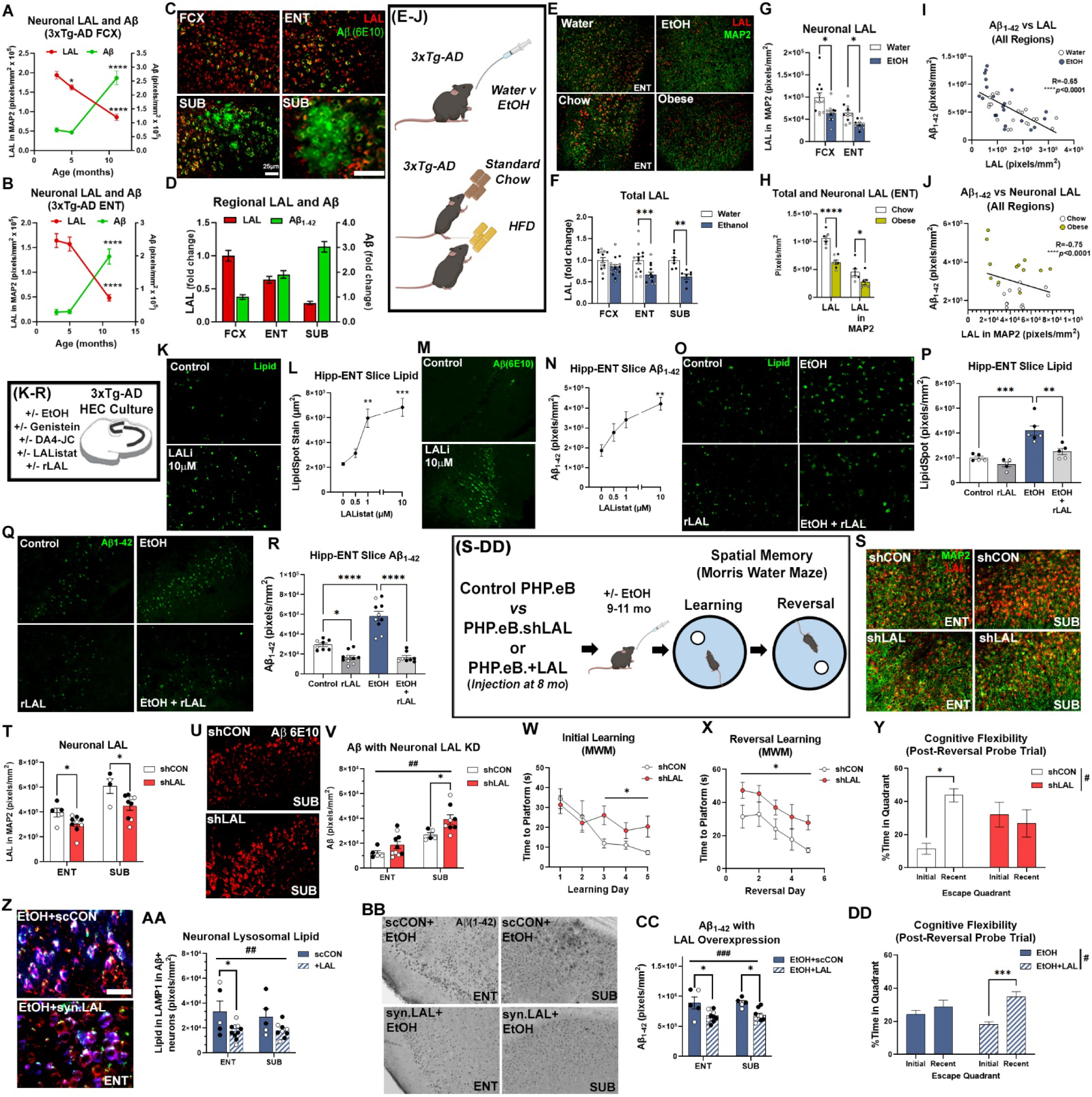
Loss of neuronal LAL drives Aβ accumulation. **(A, B)** Loss of neuronal LAL precedes Aβ accumulation in AD mouse FCX and ENT. **(C, D)** 3xTg-AD regions where FCX had the highest levels of LAL and the lowest levels of Aβ, ENT had intermediate levels of both, and subiculum (SUB) had Aβ plaques and low levels of LAL. **(E-H)** Both ethanol and obesity caused a loss of total and neuronal LAL. **(I, J)** Strong negative correlations were seen between Aβ_1-42_ and LAL in all subjects and regions with both ethanol and obesity. **(K-R)** *Ex vivo* studies using hippocampal-entorhinal brain slice cultures (HEBSC) from adult 3xTg-AD mice found a causal relationship between LAL and Aβ. LAL inhibitor LAListat (LALi) caused a dose-dependent increase in **(K, L)** lipid and **(M, N)** intraneuronal Aβ in ENT region of HEBSC. Recombinant LAL blocked ethanol-induced increases in **(O, P)** lipid and **(Q, R)** intraneuronal Aβ. \*\**p*<0.01, \*\*\**p*<0.0001, one-way ANOVA, Tukey’s multiple comparisons test. **(S, T)** Neuronal expression of shRNAs against LAL in 3xTg-AD reduced neuronal LAL in ENT and SUB (F_1,23_=23.98, *p*<0.001). Neuronal LAL knock-down **(U, V)** increased intraneuronal Aβ (F_1,24_=8.3, ##*p*<0.01, \**p*<0.05, Sidak’s post-test) and impaired **(W)** initial learning in the Morris water maze (MWM, F_1,13_=5.72, \**p*<0.05, repeated measures ANOVA), **(X)** reversal learning (F_1,11_=5.2, \**p*<0.05), and **(Y)** cognitive flexibility (F_1,24_=5.77, treatment x quadrant, #*p*<0.05, \**p*<0.05, Sidak’s). Neuronal overexpression of LAL in 3xTg-AD prior to ethanol treatment **(Z, AA)** reduced NLL (F_1,25_=9.5, ##*p*<0.01, \**p*<0.05, Sidak’s) **(BB, CC)** lowered Aβ_1-42_ in ENT and SUB (F_1,25_=18.52, ###*p*<0.001, \**p*<0.05, Sidak’s) and **(DD)** improved cognitive flexibility deficits (F_1,24_=4.39, treatment x quadrant #*p*<0.05, \*\*\**p*<0.001, Sidak’s).

Next, to determine the impact of neuronal LAL loss on Aβ pathology *in vivo*, 8 months old 3xTg-AD mice received either *PHP.eB.syn.shLAL* or *PHP.eB.shCON* i.v. injections. As expected, *PHP.eB.syn.shLAL* reduced neuronal LAL in the ENT and SUB (Figure 5S-T), causing increases in total lipid (Extended Data Figure 5E-F), neuronal lipid (Extended Data Figure 5G-H), and Aβ (Figure 5U-V). Accordingly, the level of Aβ was strongly correlated with neuronal lipid within Aβ+ neurons across regions (Extended Data Figure 5I, R=0.69, \*\**p*<0.0001). Mice with neuronal LAL knockdown showed deficits in learning in the Morris water maze (Figure 5W) as well as impaired reversal learning and cognitive flexibility consistent with a worsening of AD progression (Figure 5X-Y). To determine if LAL overexpression could improve ethanol-induced enhancement of AD pathology, mice received *PHP.eB.syn.WPRE.LAL* or *PHP.eB.scCON* injection at 8 months, followed by chronic ethanol. *PHP.eB.syn.WPRE.LAL* increased overexpression of *LAL* in 3xTg-AD cortex and hippocampus (Extended Data Figure 5J), where it reduced cytosolic lipid (Extended Data Figure 5K-L), NLL (Figure 5Z-AA), and intraneuronal Aβ in ENT and subiculum (Figure 5BB-CC). This was associated with an improvement in cognitive flexibility (Figure 5DD). Together, this indicates that the loss of neuronal LAL promotes Aβ accumulation and cognitive decline *in vivo*.

**Extended Data Figure 5.**
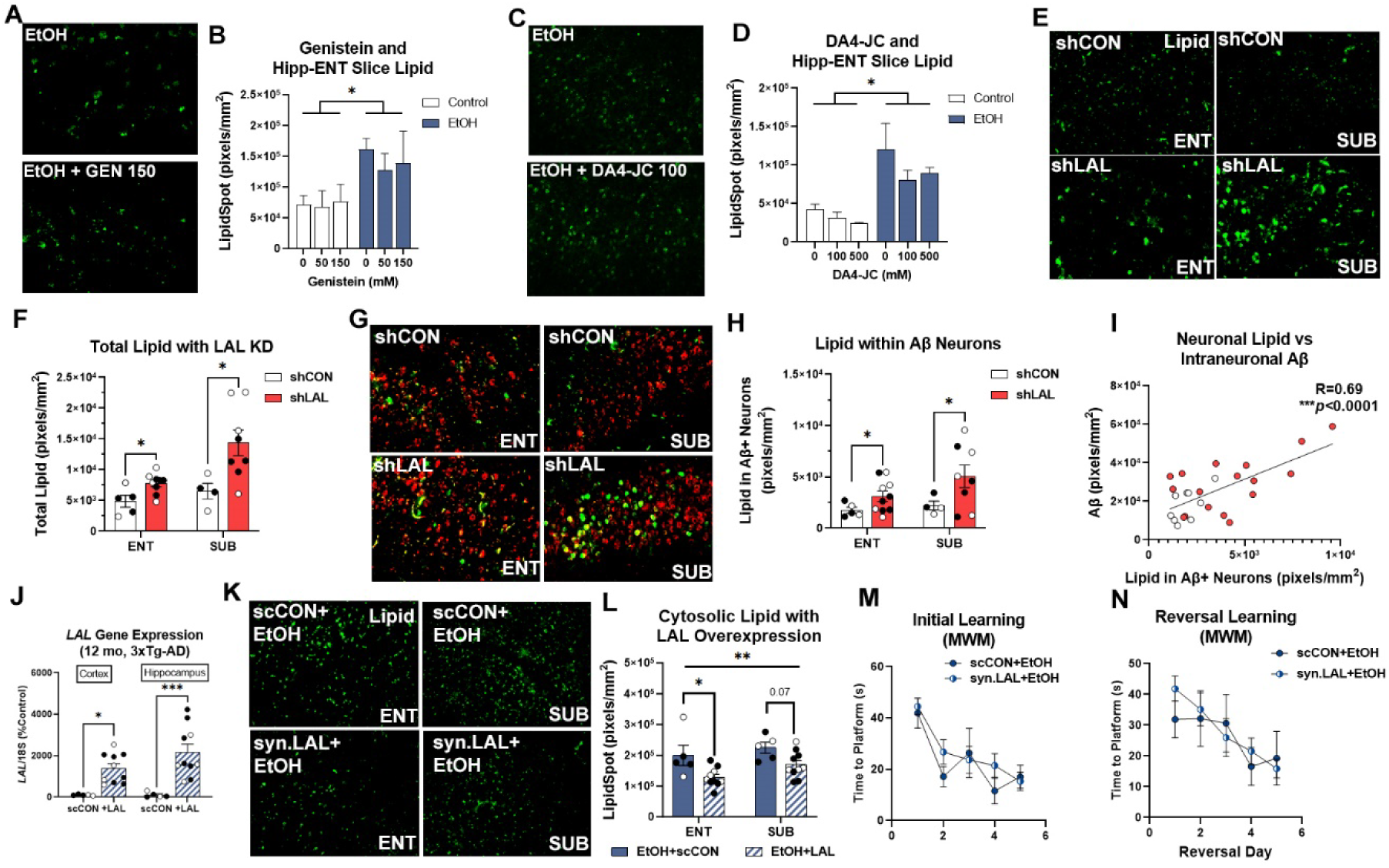
Modulation of lipid accumulation by TFEB, GLP/GIP, and LAL. **(A, B)** The TFEB agonist genistein had no effect on ethanol-induction of lipid in 3xTg-AD HEBSC. **(C, D)** GLP/GIP agonist DA4-JC had minimal effect on lipid induced by ethanol in 3xTg-AD HEBSC. PHP.eB.syn.shLAL viruses increased **(E, F)** total cytosolic lipid and **(G, H)** neuronal lipid in ENT and SUB of 3xTg-AD mice. **(I)** A strong positive correlation was found between intraneuronal Aβ and cytosolic lipid levels was found within Aβ-containing neurons. **(J)** PHP.eB.syn.WPRE.LAL viruses increased LAL gene expression in cortex and hippocampus. **(K, L)** Neuronal LAL overexpression prior to ethanol treatment reduced cytosolic lipid levels in ENT and SUB of 3xTg-AD mice. **(M, N)** Neuronal LAL overexpression had no effect on learning or reversal in the Morris water maze (MWM).

### LAL loss in healthy human aging brain and LOAD with elongating polymerase pausing

To determine if LAL loss translates to humans LOAD and extends beyond obesity and alcohol use, we measured LAL in brain from LOAD subjects and age-matched healthy control donors (HC) with no history of alcohol use disorder. LAL was found in neurons across regions (Figure 6A, top). However, in human LOAD, LAL levels were much lower and clear neuronal morphology was lost (Figure 6A, bottom). Robust reductions in LAL were found in the ENT (Figure 6B, 47%,), CA (Figure 6C, 70%), subiculum (Figure 6D, 56%), dentate (Figure 5E, 27%), and the vm-PFC (Figure 6F, 62%). In LOAD, Aβ_1-42_ was increased in these regions (Extended Data Figure 6A-H) with strong negative correlations between LAL and Aβ_1-42_ across all subjects in CA1 (Figure 6G, R= -0.60, \*\*\*\**p*<0.0001) and ENT (Figure 6H, R= -0.5, \*\**p*<0.0015). Similar to mice, LAL protein declined with age in healthy subjects in CA1 (Figure 6I, R= -0.55, \*\**p*<0.004) and ENT (Figure 6J, R= -0.55, \*\**p*<0.01). Expression of the LAL gene (*LIPA)* was reduced in LOAD (Figure 6K) with promoter occupancy of its primary transcription factor, FOXO1, greatly increased, though total FOXO1 levels were unchanged (Figure 6L, Extended Data Figure 6I). Binding of the active RNA polymerase II (p-RBP1) was increased in from promoter region to exon 3, with a reduced localization compared to HCs at later exons (Figure 6N). Further, in HCs, *LIPA* gene expression was positively correlated with p-RBP1 promoter occupancy (Extended Data Figure 6J, R=0.64, \**p*<0.05), while in LOAD, no association between p-RBP1 promoter occupancy and gene expression was found (Extended Data Figure 6K). These findings are consistent with pausing of elongating RNA polymerase, resulting in reduced LAL transcription and subsequent promotion of Aβ pathology.

**Figure 6.**
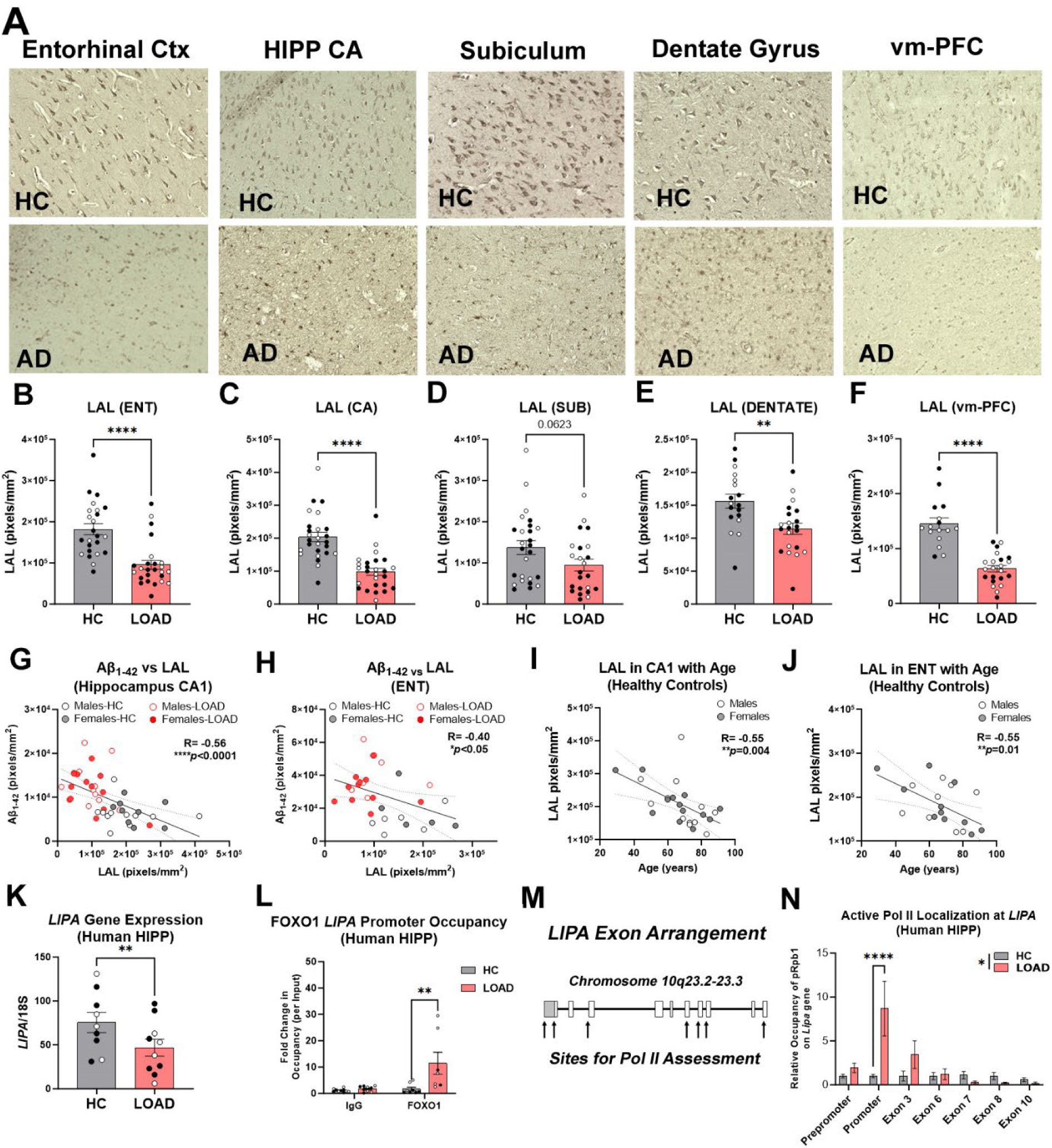
Loss of LAL in humans with LOAD and across normal aging with polymerase pausing. **(A)** LAL IHC finding a profound loss of LAL in human LOAD. In healthy control (HC) subjects, LAL was seen in cells with neuronal morphology, which was reduced in LOAD. **(B)** 47% loss of LAL in ENT, **(C)** 52% loss in CA1, **(D)** trend toward a 31% reduction in subiculum, a **(E)** 27% loss in the dentate gyrus, and a **(F)** 56% loss on the vm-PFC. paired *t-*test \*\*\*\**p*<0.0001, \*\**p*<0.01. Open circles-males filled circles-females. Strong negative correlations between LAL and Aβ_1-42_ staining were seen across all subjects in **(G)** CA1 (R= -0.55, \*\*\*\**p*<0.0001 and **(H)** ENT (R= -0.4, \**p=*0.05). In healthy control subjects, LAL declined with age in **(I)** CA1 and **(J)** ENT. **(K)** RT-PCR found a reduction in expression of the LAL gene *LIPA.* **(L)** FOXO1 CHIP found increased *LIPA* promoter occupancy in LOAD. **(M)** *LIPA* gene arrangement with arrows denoting sites assessed for Poll II localization. **(N)** Active Poll II (pRpb1) CHIP found increased Poll II association with the *LIPA* promoter region and early exon 3 in LOAD with reduced localization at later exons 7 and 8. 2-way *ANOVA* LOAD main effect F_1,68_ = 5.062, *p=*0.028. \*\**p*<0.01, Sidak’s post-test.

**Extended Data Figure 6.**
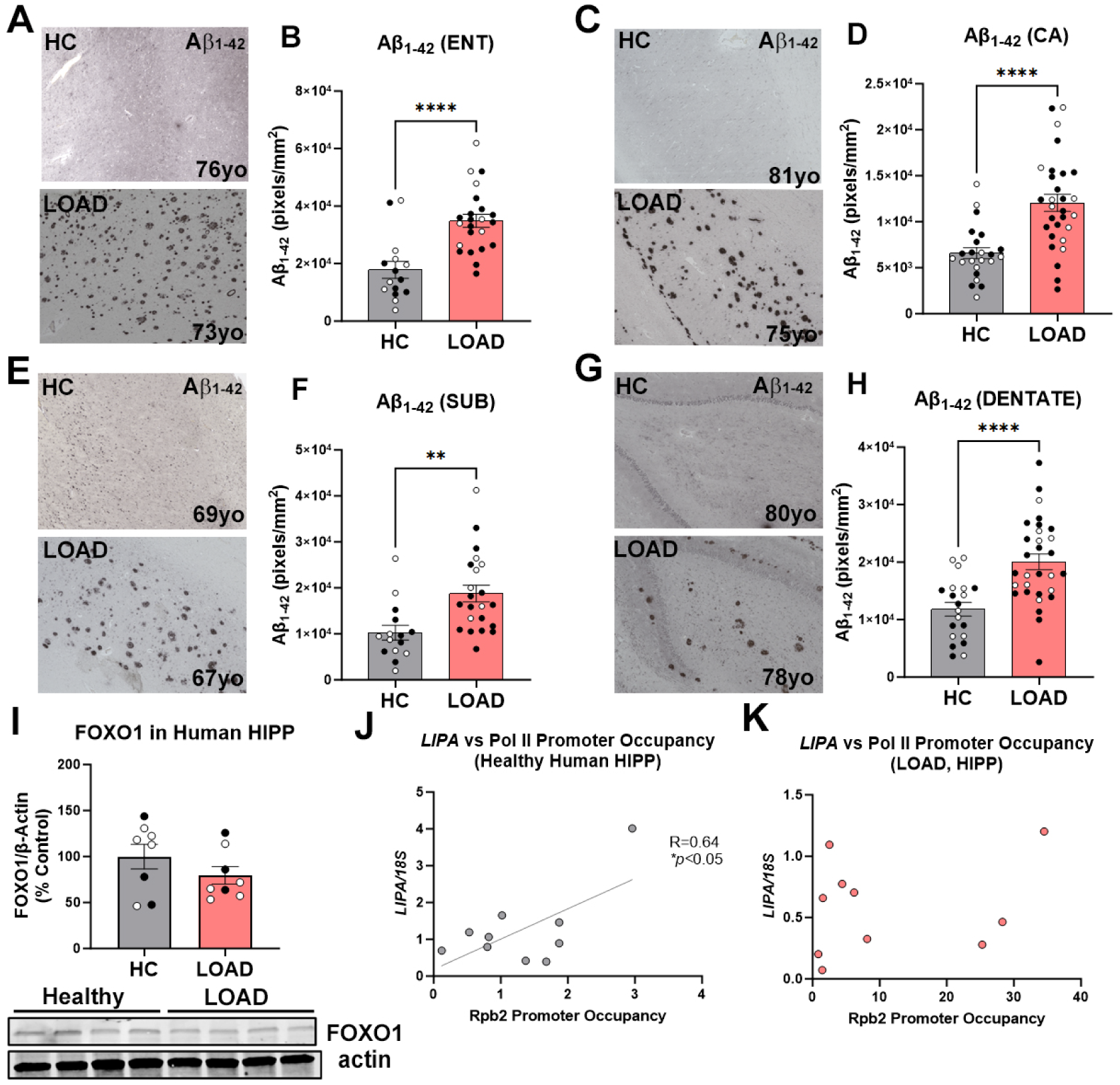
Human LOAD pathology. Healthy control (HC) and tissue from subjects with LOAD were assessed for AD amyloid pathology. Aβ_1-42_ staining was increased in the **(A, B)** ENT, **(C, D)** CA regions of hippocampus, **(E, F)** SUB, and **(G, H)** dentate gyrus. **(I)** FOXO1 protein levels were unchanged in LOAD by Western blot. **(J)** A strong positive correlation was found between *LIPA* gene expression and Rpb2 promoter occupancy in the hippocampus of HC subjects but not in **(K)** individuals with LOAD.

## Concluding Remarks

There is ongoing debate regarding early drivers of LOAD pathogenesis. The emergence of intraneuronal Aβ followed by extracellular plaque formation is subsequently followed by tau pathology^10^. However, the underlying cellular mechanisms that cross multiple LOAD behavioral risk factors and cause amyloid accumulation have not been defined. Identification of these molecular changes are essential for early diagnostic and prevention efforts. GWAS and transcriptomic studies have suggested roles for lipid metabolism in AD pathogenesis, though the cellular mechanisms that play a role in AD pathogenesis have been unclear^6–9,35,36^. By comparing two distinct midlife LOAD risk factors, we found that the loss of neuronal LAL precedes and promotes Aβ pathology and cognitive deficits by promoting the accumulation of lipid in neuronal lysosomes (Extended Data Figure 7). Evidence for this was found *in vivo* (AD and WT mice), *ex vivo,* and in human LOAD brain. Increased NLL reduced localization of Aβ to lysosomes, which may include both disruption in endolysosomal fusion and reduced lysosomal efficiency, as supported by reduced lysotracker and downregulation of vATPases. This would be consistent with recent work finding a reduction of autolysosomal acidification precedes intraneuronal Aβ accumulation and subsequent plaque formation^18^. Interestingly, genistein, an agonist of the master lysosomal transcription factor TFEB, did not reduce lipid accumulation, suggesting that normalizing lysosomal acidification alone may not be sufficient to prevent disease progression. However, LAL supplementation normalized both lysosomal lipid and Aβ both *in vitro* and *in vivo*. This suggests that normalizing lysosomal acidification alone may not be sufficient to prevent disease progression. Neuronal knock-down of LAL in AD mice increased Aβ accumulation and caused deficits in memory and cognitive flexibility, while neuronal LAL overexpression blunted increases in Aβ pathology and improved cognitive flexibility. The loss of LAL in human postmortem LOAD subjects, who did not have significant alcohol use or obesity, suggests this is a fundamental feature of LOAD pathogenesis that extends beyond these two risk factors.

**Extended Data Figure 7.**
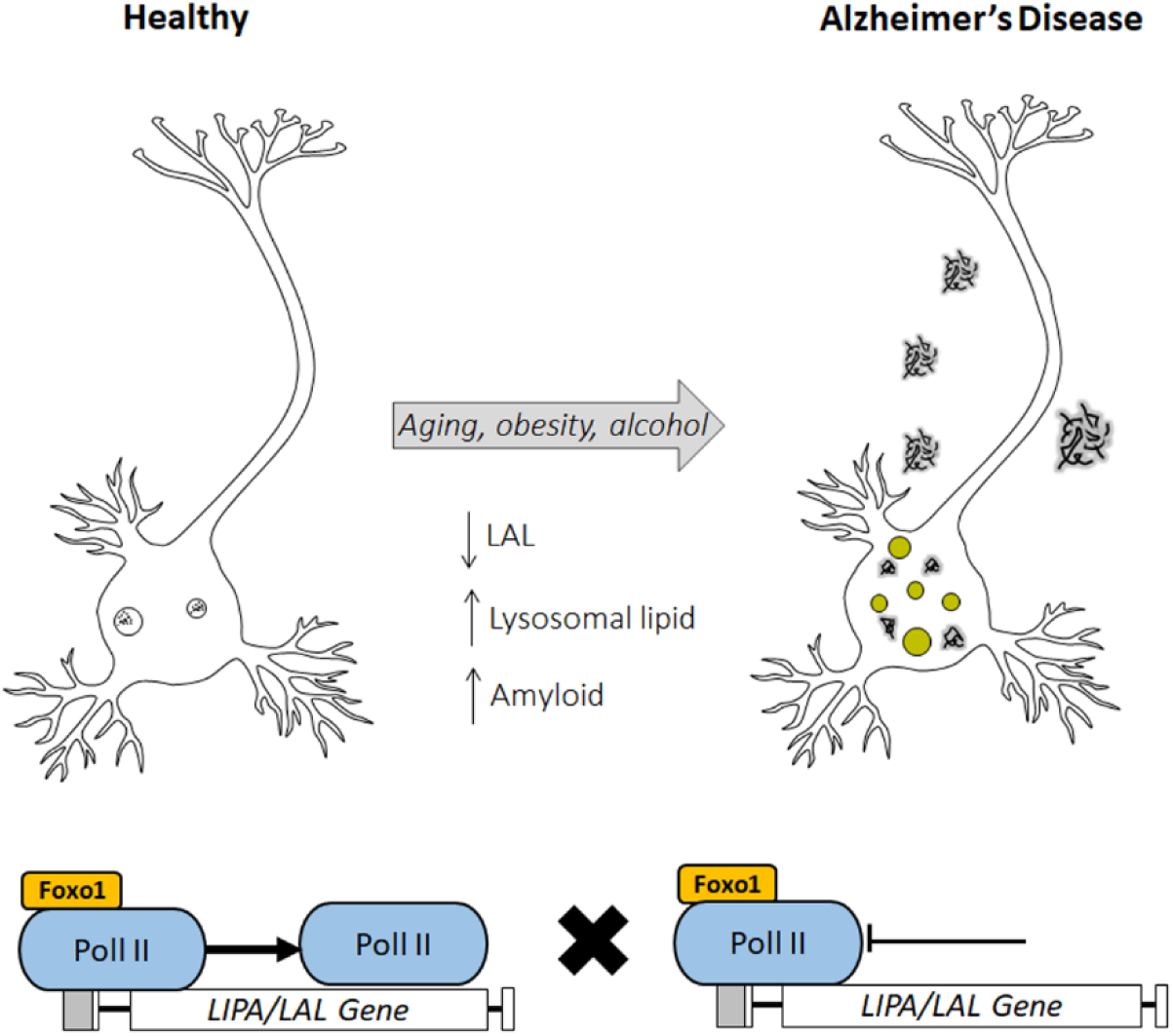
Loss of LAL promotes LOAD pathogenesis.

Aging is requisite for the development of LOAD. Though the 3xTg-AD model features lifelong expression of human familial AD transgenes, pathology emerges slowly with aging. This work finds that the loss of LAL represents the loss of a key resilience mechanism against Aβ accumulation. This was supported by findings in WT mice and healthy human aging. NLL increased greatly with age along with an age-related loss in LAL and corresponding amyloid accumulation in the ENT, which mirrors findings in healthy human brain. This suggests that LAL loss with a subsequent increase in NLL is an age-related phenomenon that produces a neuro-environment that is vulnerable to LOAD. Work in human LOAD hippocampus found polymerase pausing at the *LAL* gene which was associated with reduced *LAL* transcription. To our knowledge this is the first report of RNA polymerase pausing mediating LOAD pathology and is consistent with recent findings of Pol II insufficiency the aged mouse liver^37^. Removing the brake on the transcription of LAL and other genes could serve as a future therapeutic approach, which is a focus of ongoing studies. Further, microglia become increasingly polarized to proinflammatory state with aging and LOAD^31,38^. Our finding that chemogenetic inhibition of microglia prevents increases in NLL implicates microglia as regulators of neuronal lipid metabolism and suggests aged microglia can promote LOAD pathology by inducing neuronal lipid changes. Future studies will investigate the mechanisms by which microglia regulate neuronal lipid metabolism. Together, this work finds an age-related loss in neuronal LAL in combination with midlife LOAD risk factors, or perhaps other lipid-related LOAD genetic risks, results in the emergence of LOAD pathology.

In addition to reducing LAL, both alcohol and obesity decreased the expression of lipid efflux transporters (*ABCG1* and *ABCA1*) which could also promote intracellular lipid accumulation. Polymorphisms in *ABCA1* and other lipid efflux transporters such as *ABCA2* and *ABCA7* are associated with increased risk for LOAD^8,9^. Further, both alcohol and obesity altered expression of lipid coating *PLIN* proteins in a manner consistent with reduced lysosomal degradation of lipids. Expression of *PLIN4*, which can prevent lysosomal degradation of cytosolic lipid droplets^39^, was increased with age in WT mice and by both ethanol and obesity in 3xTg-AD mice, whereas *PLIN2* and *PLIN3*, which promote lysosomal degradation^40^, were decreased. Together, this suggests that reductions in lipid efflux in addition to lipophagy may result in NLL and reduced degradation of intraneuronal amyloid.

In summary, this work finds that the loss of neuronal LAL is a fundamental feature of LOAD pathogenesis. Further, it suggests that approaches that increase neuronal LAL or lower the intracellular neuronal lipid burden could have preventative or therapeutic value for LOAD.

## Methods

### Postmortem Human Brain Samples

Brains from human donors were from the New South Wales Brain Tissue Resource Centre (NSW-BTRC) and the Victoria Brain Bank (VBB) in Australia. Human samples were obtained under Ethics Committee Approval Number X11-0107^1,2^. Subject demographics are in Extended Data Table 1.

### Mouse Models

3xTg-AD (human APPSwe, tauP301, and Psen1^tm1Mpm^)^3,4^ male and female breeders were obtained from the Jackson Laboratory Mutant Mouse Resource & Research Centers (MMRC). WT mice were bred in house or obtained from the NIA aged mouse colony. Mice were bred and pups weaned at 30 days of age and group-housed with same-sex littermates. Animal protocols were approved by the University of North Carolina at Chapel Hill Institutional Animal Care and Use Committee (IACUC) and were in accordance with NIH regulations (Protocols 20-232.0 and 21-052.0).

### Chronic alcohol treatment

3xTg-AD or WT mice, mice received a single daily intragastric (i.g.) administration of either alcohol (i.e., ethanol) or water (five-days on/two-days off) to mimic human intermittent drinking patterns during midlife (9 months of age) for 5 to 8 weeks. Mice metabolize ethanol ∼8X faster than humans ^5^. Therefore, a dose was given that produces an average blood alcohol concentration (BAC) of ∼0.1 mg/dL during the 12 hours of intoxication (5g/kg/d, i.g., 20% ethanol w/v, peak BAC ∼280mg/dL at 1 hour) ^6,7^. No differences in body weight were found between treatment groups. Mice were sacrificed 24h after final administration and tissue was collected for tissue analyses.

### Western diet-induced midlife obesity

3xTg-AD or WT mice were fed either a control complete diet (3.6kcal/g, 14.8% fat) or a Western diet to induce obesity (4.5 kcal/g, 21.2% fat) from 6 to 11 months of age. Mice were sacrificed and tissue was collected for analysis.

### Microglial Gi DREADD inhibition

For *ex-vivo* slice culture experiments HEBSCs from 3xTg-AD mice were transfected with hM4di into microglia (AAV9.CD68.hM4di for 24h) as previously reported^8,9^ and then treated with ethanol (100mM, 4 days) +/- CNO (1µM). For *in vivo* studies, heterozygous CX3CR1.Cre^ERT2^.hM4di were treated with tamoxifen (75 mg/kg/d, i.p.) for 5 days during adulthood (12 weeks of age). After 4 weeks to allow for repopulation of peripheral monocytes, mice received either ethanol (5g/kg/d, i.g.) or water for 10 days +/- CNO (3mg/kg, 10 hours after ethanol) and sacrificed 24 hours after the last ethanol treatment.

### Perfusion and tissue collection for immunohistochemistry

At the conclusion of each experiment, subjects were sacrificed by transcardial perfusion with 0.1 M phosphate-buffered saline (PBS, pH 7.4), and brains excised and hemisected. On one hemisphere, the cortex and hippocampus were dissected and snap frozen in liquid nitrogen for protein and RNA analyses as we have reported previously^10^. The other hemisphere was drop-fixed in 4.0% paraformaldehyde for immunohistochemical assessments. Coronal sections were cut (40 μm) on a sliding microtome (MICROM HM450; ThermoScientific, Austin, TX), and sections were sequentially collected into well plates and stored at -20°C in cryoprotectant (30% glycol/30% ethylene glycol in PBS).

### RT-PCR

mRNA was extracted from frozen cortex or hippocampus as reported^10^. Briefly, samples were homogenized with Trizol (Invitrogen) and RNA was isolated by chloroform extraction, followed by reverse transcription as described previously^11^. SYBR green PCR master mix (Applied Biosystems, Foster City, CA) was used for qRT-PCR analysis. Primer sequences were designed using the National Library of Medicine Primer-BLAST tool or were obtained from PrimerBank database (Extended Data Table 2)^12–14^. Only primers with no predicted non-specific targets and single peak melt curves were used. Genes of interest were normalized to the expression of the reference gene 18S using the cycle threshold (Ct) value of each target gene product. The ΔΔCt method was used to compare relative differences between control and treatment groups, and the ratio by 18S or the percent change relative to 18S were used in analysis.

### Western Blot

Western blot was performed as we have reported previously^9,10^. Brain tissue from the cerebral cortex was homogenized in lysis buffer (Tris-HCl, pH 7.5, Sucrose, EDTA, EGTA, 1% Triton X-100, protease, and phosphatase inhibitors). Forty milligrams of protein were loaded into each lane on SDS polyacrylamide gels and were transferred to PVDF membranes. Membranes were washed in TBS and blocked for 1 hour at room temperature (Li-Cor Blocking Solution; 92760001) then were incubated overnight at 4° Celsius with the primary antibodies listed in Extended Data Table 2. Membranes were washed in TBS with 0.1% Tween-20 (Sigma-Aldrich, St. Louis, MO) then were incubated in the appropriate conjugated secondary antibody (Rockland H&L Pre-absorbed). Membranes were washed again in TBS and visualized using LiCor Image Studio Lite Ver 5.2. Western Blots were analyzed using Image Studio Lite software and each protein of interest was normalized to housekeeping protein GAPDH. GAPDH expression was not affected by ethanol treatment or high-fat diet. The protein of interest was normalized to GAPDH for each sample and the percentage change relative to controls was calculated for each blot.

### Immunohistochemistry (IHC) and Immunofluorescence (IF)

Free-floating sections (40µm) were washed in 0.1M PBS, quenched in 0.6% H_2_O_2_ for 30 minutes to inhibit endogenous peroxidases, then incubated at 70°C in pH=6.0 1X Citrate Buffer for antigen retrieval. To allow membrane permeabilization, sections were blocked for 1 hour at room temperature in 4% normal serum with 0.1% Triton X-100. Sections were incubated in the relevant primary antibody (please see above) in blocking solution at 4°C overnight. Negative controls for non-specific binding were done using the same protocol, omitting the primary antibody. The next day sections were incubated at room temperature for one hour with a biotinylated secondary antibody (1:200, Vector Laboratories, Burlingame, CA), washed briefly in PBS, then incubated for 1 hour in avidin-biotin complex (ABC) (Vectastain ABC Kit; Vector Laboratories). Nickel-enhanced diaminobenzidine (Sigma-Aldrich) was used as the chromogen for visualization. Sections were mounted on charged glass slides, allowed to dry, dehydrated in a series of ethanol, and covered using Cytoseal (Fischer). For IF, the day after primary antibody incubation, sections were washed with PBS and incubated for 1 hour at room temperature with the respective Alexa Fluor conjugated secondary antibodies (1:1000; Invitrogen, Carlsbad, Ca, USA). If staining for neutral lipid droplets was being performed, sections were washed with PBS then stained with 1xLipidSpot AlexaFluor 488 (Biotium, San Francisco, CA) in PBS for 20 min. Sections were mounted and cover slips placed with Prolong Gold Anti-Fade mounting medium with DAPI (Invitrogen; Carlsbad, CA, USA). Negative control for non-specific binding was done separately with the exception that the primary antibody was omitted.

The Keyence BZ-X800 all-in-one Microscope was used for representative images and microscopic analysis of tissue. For each region of interest assessed, images were taken on 3-5 identical sections per subject at a representative location, selected using the mouse brain atlas^15^. Bregma used by region were: entorhinal cortex and subiculum (-3.52 to -4.04mm), and CA1 (-2.92 to -3.40mm). The immunoreactivity (+IR) was quantified in each region using either the BZ-X800 software or ImageJ Analysis Software and reported as +IR pixels/mm^2^.

### Hippocampal-Entorhinal Brain Slice Culture (HEBSC)

HEBSC was performed using techniques routinely used in our laboratory ^2,9,11^. Briefly, sections were prepared from either the P7 or adult (6 mo) 3x-Tg-AD hippocampal-entorhinal cortex formation as described by Stoppini et al^16^. Initial experiments were done on HEBSC from P7 mice then in HEBSC from adults to ensure developmental differences did not drive findings. Mice were decapitated, brains extracted and the hippocampal-enthorhinal complex dissected in Gey’s bugger (Sigma-Aldrich). Slices were cut transversely at 375 µm using a McIlwain tissue chopper and placed on Millicell culture inserts (Millipore, PICMORG50, up to 13 slices per insert). Slices acclimated in culture media (MEM plus 24mM HEPES and Hank’s salts, 25% horse serum, 5.5 g/L glucose, 2mM L-glutamine) in a humified 5% CO2 incubator for seven days, followed by 12% horse serum for four days, and 6% horse serum until the completion of the experiments.

### Behavior

#### Open field

Exploratory activity in a novel environment was used to insure none of the mice had overt motor impairment or were notably hypoactive (a sign of possible health issues) before starting evaluation in the Morris water maze. This assessed by a one-hour trial in an open field chamber (41 cm x 41 cm x 30 cm) crossed by a grid of photobeams (VersaMax system, AccuScan Instruments). Counts were taken of the number of photobeams broken during the trial in 5-min intervals, with separate measures for locomotor activity (total distance traveled) and vertical rearing movements. Time spent in the center region was used as an index of anxiety-like behavior.

#### Morris water maze

The water maze was used to evaluate spatial and reversal learning, swimming ability, and vision. The water maze consisted of a large circular pool (diameter = 122 cm) partially filled with water (45 cm deep, 24-26° C), located in a room with numerous visual cues. The procedure involved a visible platform test, acquisition in the hidden platform task, and a test for reversal learning (an index of cognitive flexibility). <u>Visible</u> <u>platform test.</u> Each mouse was given 4 trials per day, across 2 days, to swim to an escape platform cued by a patterned cylinder extending above the surface of the water. For each trial, the mouse was placed in the pool at 1 of 4 possible locations (randomly ordered), and then given 60 sec to find the visible platform. If the mouse found the platform, the trial ended, and the animal was allowed to remain 10 sec on the platform before the next trial began. If the platform was not found, the mouse was placed on the platform for 10 sec, and then given the next trial. Measures were taken of latency to find the platform and swimming speed via an automated tracking system (Noldus Ethovision). <u>Acquisition and reversal learning in a hidden platform task</u>. Following the visible platform task, mice were tested for their ability to find a submerged, hidden escape platform (diameter = 12 cm). Each mouse was given 4 trials per day, with 1 min per trial, to swim to the hidden platform. The criterion for learning was an average group latency of 15 sec or less to locate the platform. Mice were tested until the group reached criterion, with a maximum of 9 days of testing. When the group reached criterion (on day 5 in the present study), mice were given a one-min probe trial in the pool with the platform removed. Selective quadrant search was evaluated by measuring percent time in the quadrant where the platform (the target) had been placed during training, versus the opposite quadrant, and number of crosses over the target location where the platform had been placed, versus the corresponding area in the opposite quadrant. Following the acquisition phase, mice were tested for reversal learning, using the same procedure as described above. In this phase, the hidden platform was re-located to the opposite quadrant in the pool. As before, measures were taken of latency to find the platform. On day 6 of testing, the platform was removed from the pool, and the group was given a probe trial to evaluate reversal learning.

### Chromatin Immunoprecipitation (CHIP)

Chromatin immunoprecipitation was performed as previously described by our laboratory^17–20^. Briefly, postmortem human hippocampal tissue from CON and AD individuals was homogenized, cross-linked with 1.0% methanol-free formaldehyde, quenched with 1.0 M glycine, lysed with lysis buffer (1.0% [v/v] SDS, 10 mM EDTA, 50 mM Tris-HCl [pH 8.0]), and chromatin sheared to fragments of <1000 bp on a Covaris ME220. Input DNA fractions were removed from the sheared chromatin to be processed separately and the remaining sheared chromatin was incubated overnight at 4°C with an antibody against rabbit FoxO1, rabbit phospho-Rpb1, or the negative control Rabbit IgG. Protein A Dynabeads were added and rotated at 4°C for 1 hr followed by five washes in ChIP wash buffer. Both immunoprecipitated DNA and input DNA were eluted in 10% (w/v) Chelex by boiling at 95°C for 10 min followed by centrifugation. ChIP-enriched DNA was analyzed using qPCR with SSOAdvanced Universal SYBR Green Supermix (Bio-Rad, Berkeley, CA) using primers designed against regions of the *LIPA* gene. The ΔΔCt method was used to determine fold occupancy relative to control and was normalized to the input DNA fraction.

### Statistical Analyses

The specific statistical test used is noted for each assessment above and were performed in GraphPad Prism^TM^. For preplanned orthogonal contrasts, *t-*tests were used. For age-matched assessments paired *t-*tests were employed. 1-way or 2-way ANOVAs were used for multiple-group assessments. Dunnett’s or Sidak’s post-tests were used for ANOVAs when appropriate. Outliers were detected using the Grubb’s test.

## Acknowledgments

This work was supported by grants awarded to L.C. from NIH (R01AA028924, K08AA024829), R.V. (R01AG072894, K01AA025713), the Mouse Behavioral Phenotyping Core through the Carolina Institue for Developmental Disabilities (NICHD; P50HD103573; PI: Gabriel Dichter) and the Bowles Center for Alcohol Studies. This NSW-receives financial support from the National Health and Medical Research Council of Australia-Schizophrenia Research Institute as well as the National Institute of Alcohol Abuse and Alcoholism (NIAAA, R24AA012725). The senior author thanks God for His guidance and inspiration.

## Author Contributions

Conceptualization, A.M.B. and L.G.C.; Methodology, A.M.B., J.Z., V.D., S.M., S.D., C.D., R.P.V, and L.G.C.; Validation, A.M.B., L.D., E.M., M.C., H.K., K.K.; Formal Analysis, A.M.B., L.D., J.Z., V.D., S.M., C.D., R.P.V, and L.G.C., Investigation, A.M.B., L.D., J.Z., V.D., E.M., M.C., H.K., K.K., and R.P.V., Resources, J.S. and G.S., Writing-Original Draft, A.M.B., S.M., and L.G.C., Writing-Review & Editing, A.M.B., G.S., S.M., and L.G.C., Visualization, A.M.B., S.M., and L.G.C., Supervision, A.M.B., L.D., S.M., L.G.C., Project Administration, L.G.C, Funding Acquisition, S.M., R.P.V, and L.G.C.

## Data Availability

All data generated or analyzed during the current study are included in this published article (and its supplementary information files).

## Material & Correspondence

Please address correspondence and material requests to

## Extended Data Tables

**Extended Data Table 1.**
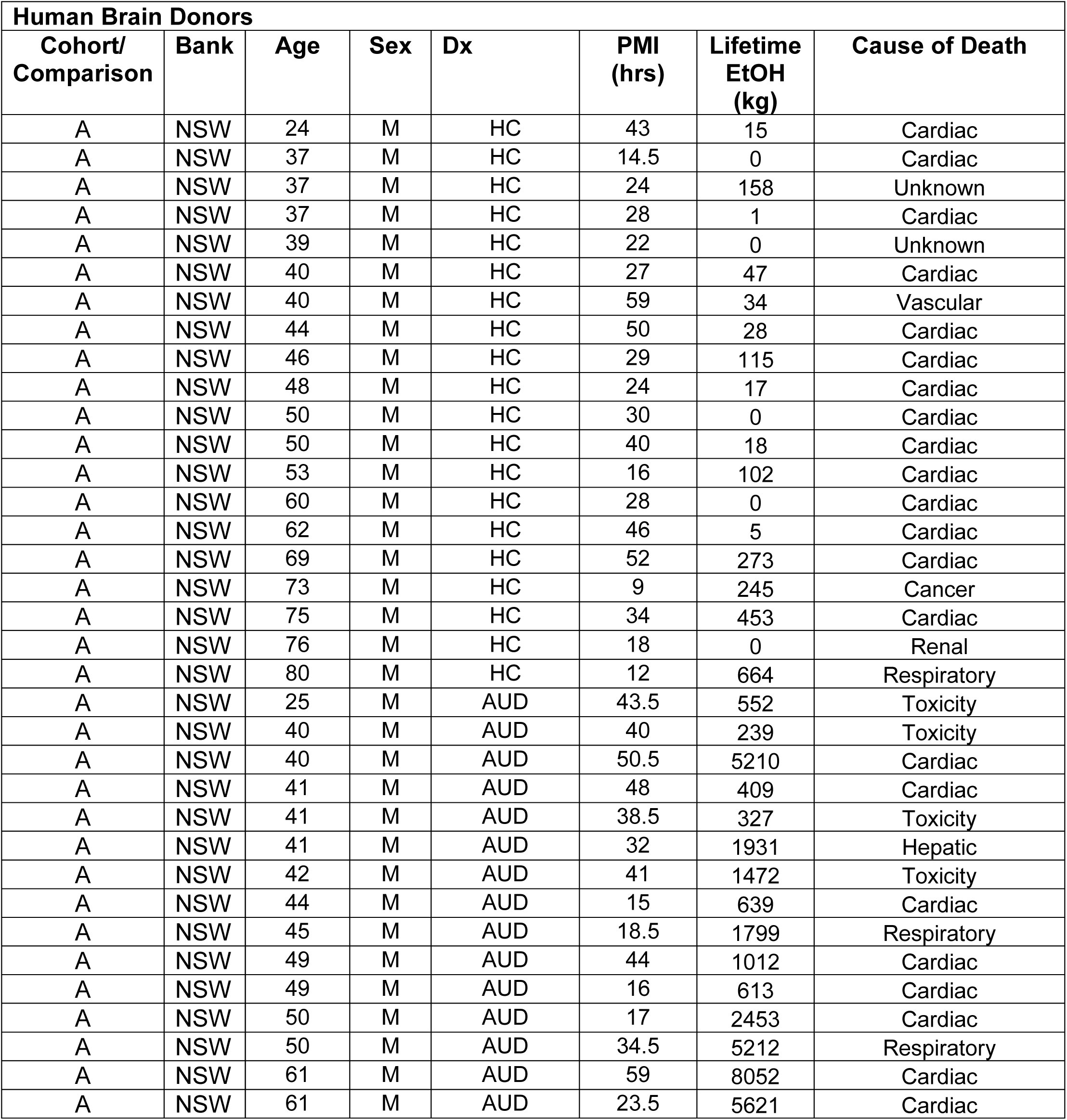
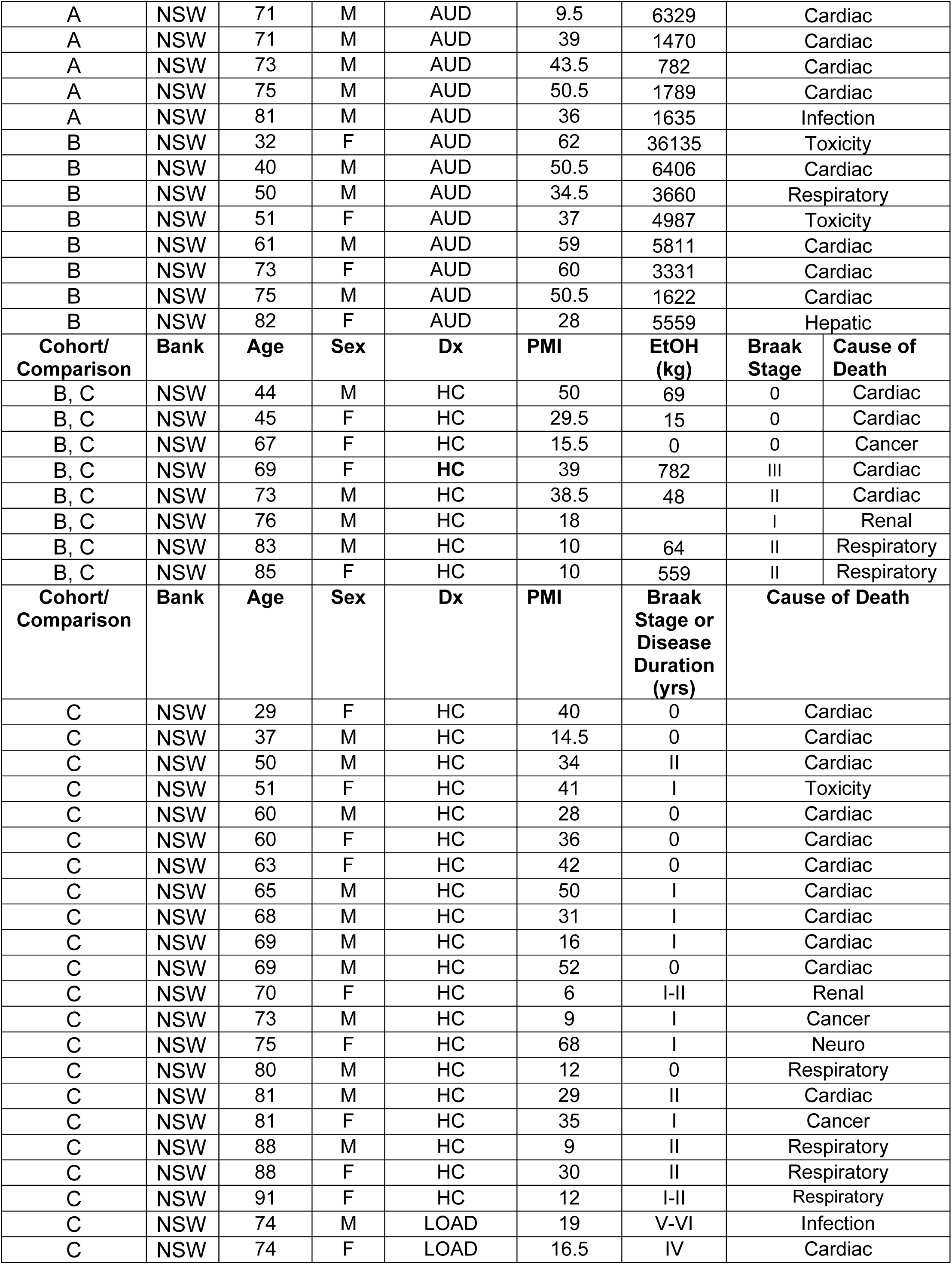
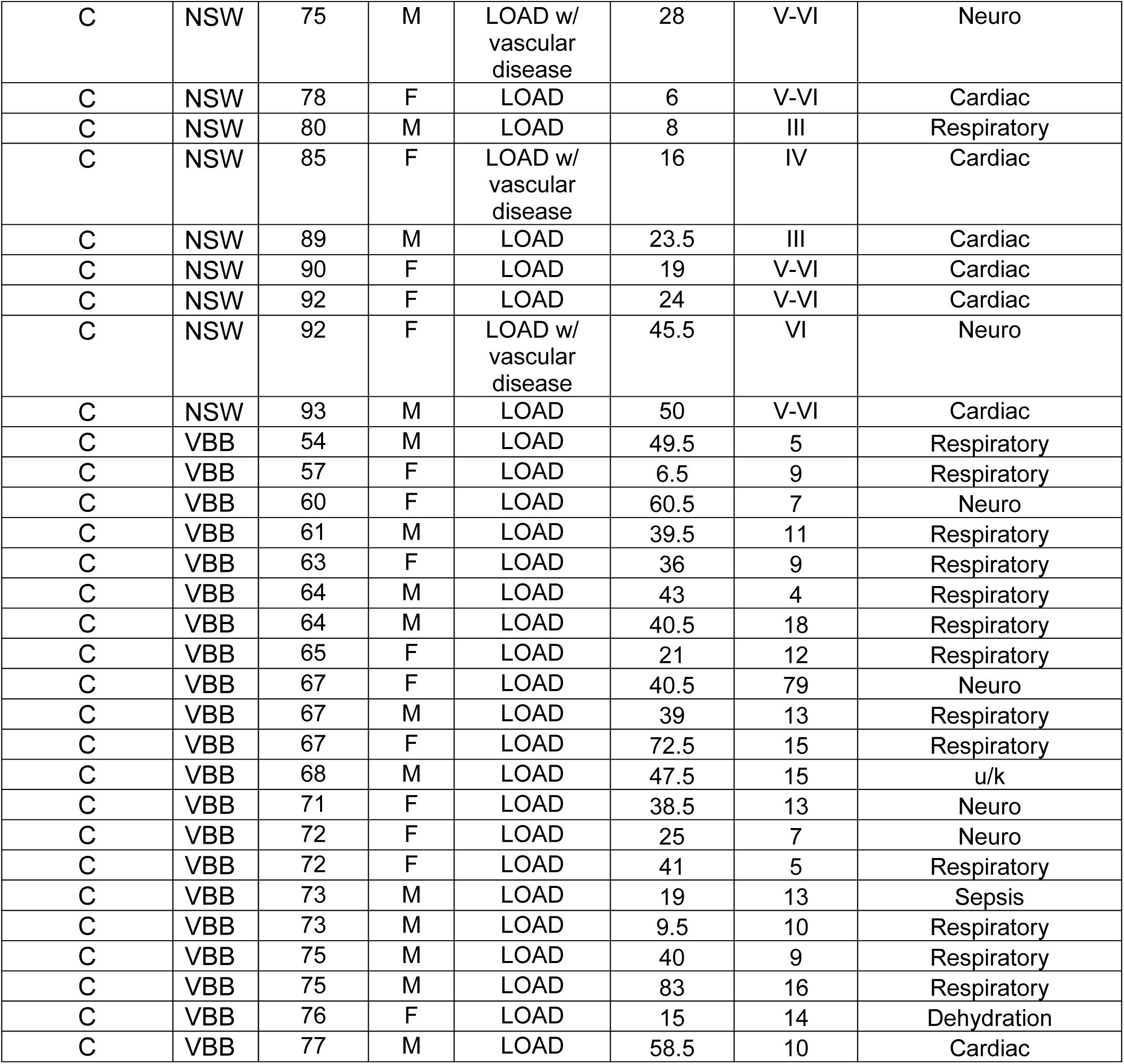
Demographics of postmortem human brain subjects. HC-Healthy controls, AUD-Alcohol Use Disorder, LOAD-Late onset Alzheimer’s disease.

| Human Brain Donors |  |  |  |  |  |  |  |
| --- | --- | --- | --- | --- | --- | --- | --- |
| Cohort/<br>Comparison | Bank | Age | Sex | Dx | PMI<br>(hrs) | Lifetime<br>EtOH<br>(kg) | Cause of Death |
| A | NSW | 24 | M | HC | 43 | 15 | Cardiac |
| A | NSW | 37 | M | HC | 14.5 | 0 | Cardiac |
| A | NSW | 37 | M | HC | 24 | 158 | Unknown |
| A | NSW | 37 | M | HC | 28 | 1 | Cardiac |
| A | NSW | 39 | M | HC | 22 | 0 | Unknown |
| A | NSW | 40 | M | HC | 27 | 47 | Cardiac |
| A | NSW | 40 | M | HC | 59 | 34 | Vascular |
| A | NSW | 44 | M | HC | 50 | 28 | Cardiac |
| A | NSW | 46 | M | HC | 29 | 115 | Cardiac |
| A | NSW | 48 | M | HC | 24 | 17 | Cardiac |
| A | NSW | 50 | M | HC | 30 | 0 | Cardiac |
| A | NSW | 50 | M | HC | 40 | 18 | Cardiac |
| A | NSW | 53 | M | HC | 16 | 102 | Cardiac |
| A | NSW | 60 | M | HC | 28 | 0 | Cardiac |
| A | NSW | 62 | M | HC | 46 | 5 | Cardiac |
| A | NSW | 69 | M | HC | 52 | 273 | Cardiac |
| A | NSW | 73 | M | HC | 9 | 245 | Cancer |
| A | NSW | 75 | M | HC | 34 | 453 | Cardiac |
| A | NSW | 76 | M | HC | 18 | 0 | Renal |
| A | NSW | 80 | M | HC | 12 | 664 | Respiratory |
| A | NSW | 25 | M | AUD | 43.5 | 552 | Toxicity |
| A | NSW | 40 | M | AUD | 40 | 239 | Toxicity |
| A | NSW | 40 | M | AUD | 50.5 | 5210 | Cardiac |
| A | NSW | 41 | M | AUD | 48 | 409 | Cardiac |
| A | NSW | 41 | M | AUD | 38.5 | 327 | Toxicity |
| A | NSW | 41 | M | AUD | 32 | 1931 | Hepatic |
| A | NSW | 42 | M | AUD | 41 | 1472 | Toxicity |
| A | NSW | 44 | M | AUD | 15 | 639 | Cardiac |
| A | NSW | 45 | M | AUD | 18.5 | 1799 | Respiratory |
| A | NSW | 49 | M | AUD | 44 | 1012 | Cardiac |
| A | NSW | 49 | M | AUD | 16 | 613 | Cardiac |
| A | NSW | 50 | M | AUD | 17 | 2453 | Cardiac |
| A | NSW | 50 | M | AUD | 34.5 | 5212 | Respiratory |
| A | NSW | 61 | M | AUD | 59 | 8052 | Cardiac |
| A | NSW | 61 | M | AUD | 23.5 | 5621 | Cardiac |

| A | NSW | 71 | M | AUD | 9.5 | 6329 | Cardiac |  |
| --- | --- | --- | --- | --- | --- | --- | --- | --- |
| A | NSW | 71 | M | AUD | 39 | 1470 | Cardiac |  |
| A | NSW | 73 | M | AUD | 43.5 | 782 | Cardiac |  |
| A | NSW | 75 | M | AUD | 50.5 | 1789 | Cardiac |  |
| A | NSW | 81 | M | AUD | 36 | 1635 | Infection |  |
| B | NSW | 32 | F | AUD | 62 | 36135 | Toxicity |  |
| B | NSW | 40 | M | AUD | 50.5 | 6406 | Cardiac |  |
| B | NSW | 50 | M | AUD | 34.5 | 3660 | Respiratory |  |
| B | NSW | 51 | F | AUD | 37 | 4987 | Toxicity |  |
| B | NSW | 61 | M | AUD | 59 | 5811 | Cardiac |  |
| B | NSW | 73 | F | AUD | 60 | 3331 | Cardiac |  |
| B | NSW | 75 | M | AUD | 50.5 | 1622 | Cardiac |  |
| B | NSW | 82 | F | AUD | 28 | 5559 | Hepatic |  |
| Cohort/<br>Comparison | Bank | Age | Sex | Dx | PMI | EtOH<br>(kg) | Braak<br>Stage | Cause of<br>Death |
| B, C | NSW | 44 | M | HC | 50 | 69 | 0 | Cardiac |
| B, C | NSW | 45 | F | HC | 29.5 | 15 | 0 | Cardiac |
| B, C | NSW | 67 | F | HC | 15.5 | 0 | 0 | Cancer |
| B, C | NSW | 69 | F | HC | 39 | 782 | III | Cardiac |
| B, C | NSW | 73 | M | HC | 38.5 | 48 | II | Cardiac |
| B, C | NSW | 76 | M | HC | 18 |  | I | Renal |
| B, C | NSW | 83 | M | HC | 10 | 64 | II | Respiratory |
| B, C | NSW | 85 | F | HC | 10 | 559 | II | Respiratory |
| Cohort/<br>Comparison | Bank | Age | Sex | Dx | PMI | Braak<br>Stage or<br>Disease<br>Duration<br>(yrs) | Cause of Death |  |
| C | NSW | 29 | F | HC | 40 | 0 | Cardiac |  |
| C | NSW | 37 | M | HC | 14.5 | 0 | Cardiac |  |
| C | NSW | 50 | M | HC | 34 | II | Cardiac |  |
| C | NSW | 51 | F | HC | 41 | I | Toxicity |  |
| C | NSW | 60 | M | HC | 28 | 0 | Cardiac |  |
| C | NSW | 60 | F | HC | 36 | 0 | Cardiac |  |
| C | NSW | 63 | F | HC | 42 | 0 | Cardiac |  |
| C | NSW | 65 | M | HC | 50 | I | Cardiac |  |
| C | NSW | 68 | M | HC | 31 | I | Cardiac |  |
| C | NSW | 69 | M | HC | 16 | I | Cardiac |  |
| C | NSW | 69 | M | HC | 52 | 0 | Cardiac |  |
| C | NSW | 70 | F | HC | 6 | I-II | Renal |  |
| C | NSW | 73 | M | HC | 9 | I | Cancer |  |
| C | NSW | 75 | F | HC | 68 | I | Neuro |  |
| C | NSW | 80 | M | HC | 12 | 0 | Respiratory |  |
| C | NSW | 81 | M | HC | 29 | II | Cardiac |  |
| C | NSW | 81 | F | HC | 35 | I | Cancer |  |
| C | NSW | 88 | M | HC | 9 | II | Respiratory |  |
| C | NSW | 88 | F | HC | 30 | II | Respiratory |  |
| C | NSW | 91 | F | HC | 12 | I-II | Respiratory |  |
| C | NSW | 74 | M | LOAD | 19 | V-VI | Infection |  |
| C | NSW | 74 | F | LOAD | 16.5 | IV | Cardiac |  |
| C | NSW | 75 | M | LOAD w/<br>vascular<br>disease | 28 | V-VI | Neuro |
| C | NSW | 78 | F | LOAD | 6 | V-VI | Cardiac |
| C | NSW | 80 | M | LOAD | 8 | III | Respiratory |
| C | NSW | 85 | F | LOAD w/<br>vascular<br>disease | 16 | IV | Cardiac |
| C | NSW | 89 | M | LOAD | 23.5 | III | Cardiac |
| C | NSW | 90 | F | LOAD | 19 | V-VI | Cardiac |
| C | NSW | 92 | F | LOAD | 24 | V-VI | Cardiac |
| C | NSW | 92 | F | LOAD w/<br>vascular<br>disease | 45.5 | VI | Neuro |
| C | NSW | 93 | M | LOAD | 50 | V-VI | Cardiac |
| C | VBB | 54 | M | LOAD | 49.5 | 5 | Respiratory |
| C | VBB | 57 | F | LOAD | 6.5 | 9 | Respiratory |
| C | VBB | 60 | F | LOAD | 60.5 | 7 | Neuro |
| C | VBB | 61 | M | LOAD | 39.5 | 11 | Respiratory |
| C | VBB | 63 | F | LOAD | 36 | 9 | Respiratory |
| C | VBB | 64 | M | LOAD | 43 | 4 | Respiratory |
| C | VBB | 64 | M | LOAD | 40.5 | 18 | Respiratory |
| C | VBB | 65 | F | LOAD | 21 | 12 | Respiratory |
| C | VBB | 67 | F | LOAD | 40.5 | 79 | Neuro |
| C | VBB | 67 | M | LOAD | 39 | 13 | Respiratory |
| C | VBB | 67 | F | LOAD | 72.5 | 15 | Respiratory |
| C | VBB | 68 | M | LOAD | 47.5 | 15 | u/k |
| C | VBB | 71 | F | LOAD | 38.5 | 13 | Neuro |
| C | VBB | 72 | F | LOAD | 25 | 7 | Neuro |
| C | VBB | 72 | F | LOAD | 41 | 5 | Respiratory |
| C | VBB | 73 | M | LOAD | 19 | 13 | Sepsis |
| C | VBB | 73 | M | LOAD | 9.5 | 10 | Respiratory |
| C | VBB | 75 | M | LOAD | 40 | 9 | Respiratory |
| C | VBB | 75 | M | LOAD | 83 | 16 | Respiratory |
| C | VBB | 76 | F | LOAD | 15 | 14 | Dehydration |
| C | VBB | 77 | M | LOAD | 58.5 | 10 | Cardiac |

**Extended Data Table 2.**
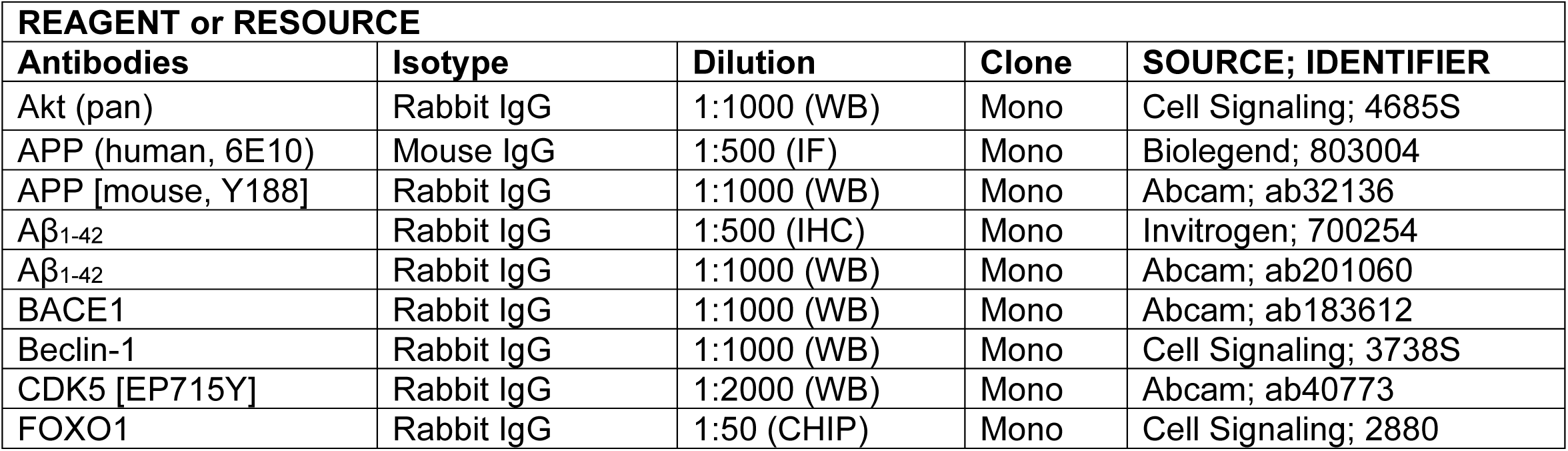
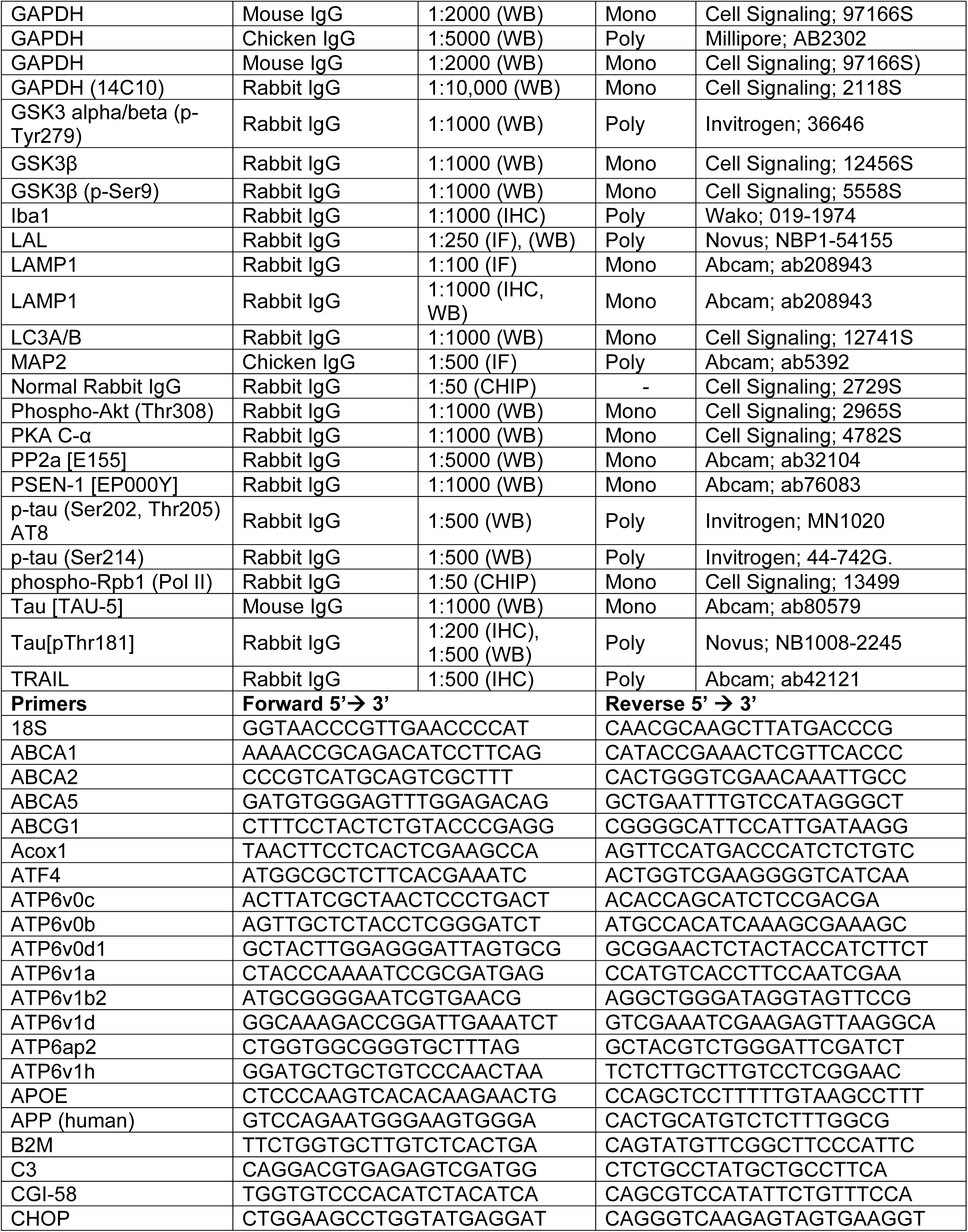
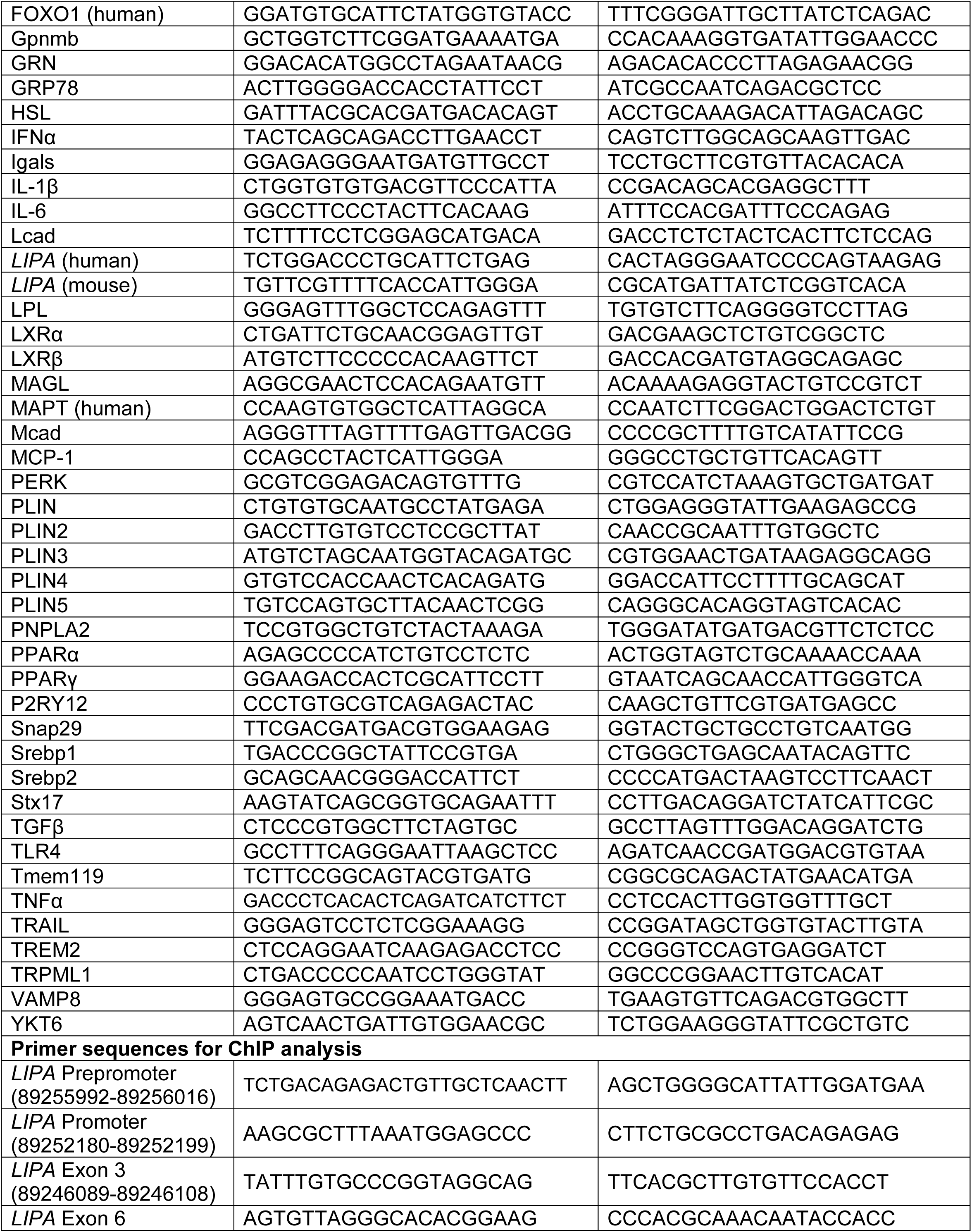
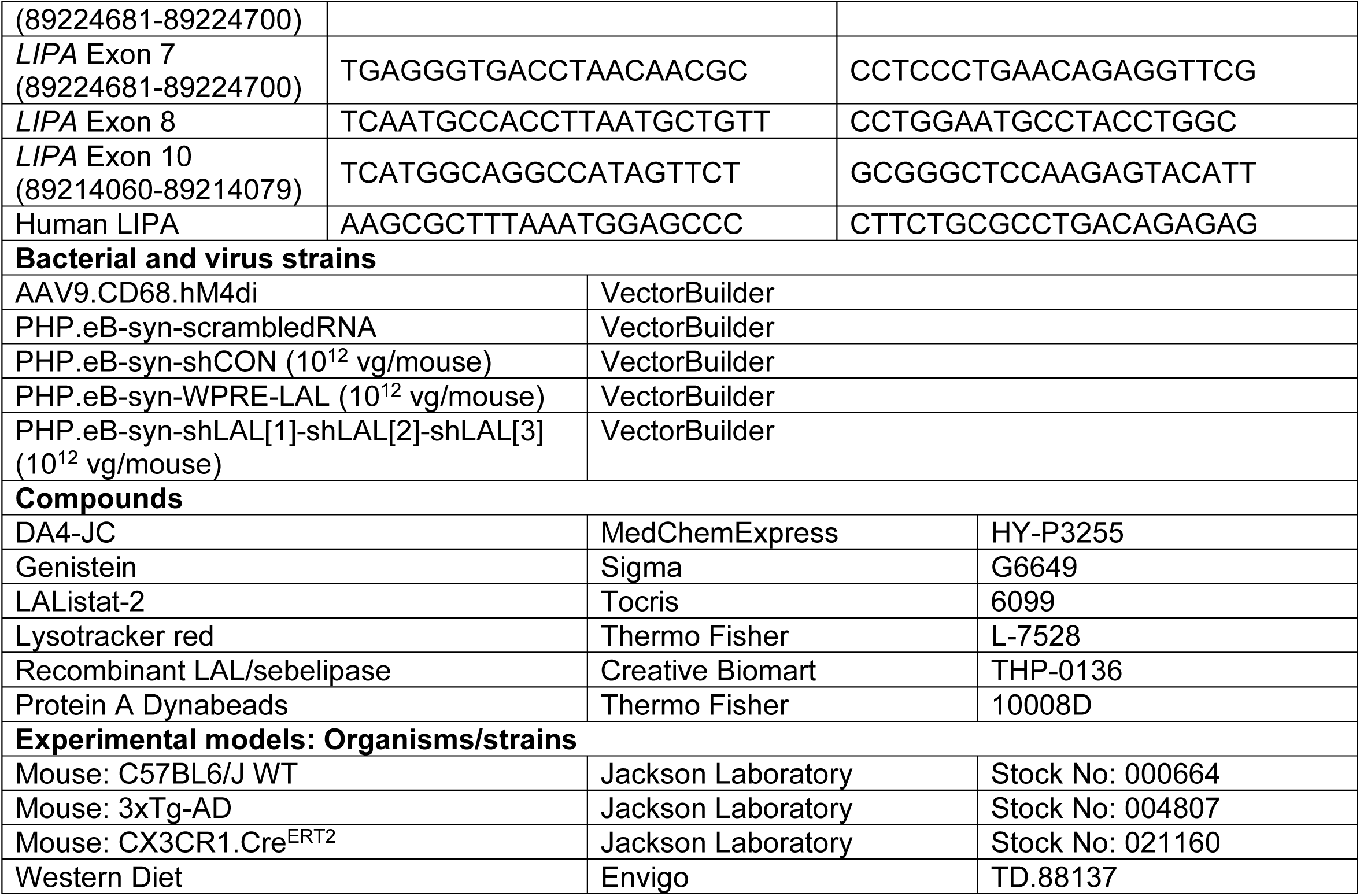
Key material resources and primer sequences.

